# Specific ABA-independent tomato transcriptome reprogramming under abiotic stress combination

**DOI:** 10.1101/2023.03.29.534684

**Authors:** Miriam Pardo-Hernández, Sara E. Martínez-Lorente, José M Martí-Guillén, Vicent Arbona, Inmaculada Simón, Rosa M Rivero

## Abstract

Crops often have to face several abiotic stresses simultaneously, and under these conditions, the plant’s response significantly differs from that observed under a single stress. Nevertheless, most of the molecular markers identified for increasing plant stress tolerance have been characterized under single abiotic stresses, explaining their unexpected results when they are tested under real field conditions. One important regulator of the plant’s responses to abiotic stresses is ABA. The ABA signaling system engages many stress-responsive genes, however, many others do not respond to ABA treatments. Thus, the ABA-independent pathway, which is still largely unknown, involve multiple signaling pathways and important molecular components necessary for the plant’s adaptation to climate change.

In the present study, tomato ABA-deficient mutants (flacca, flc) were subjected to salinity, heat, or their combination. A deep RNA-seq analysis revealed that the combination of salinity and heat induced an important reprogramming of the tomato transcriptome, and from the 685 genes that were specifically regulated under this combination in our flc mutants, 463 genes were regulated by ABA-independent systems. Among these genes, we identified 6 transcription factors (TFs) belonging to the R2R3MYB family that were significantly upregulated. A protein-protein interaction network showed that the TFs SlMYB50 and SlMYB86 were directly involved in the upregulation of the flavonol biosynthetic pathway-related genes. This is the first time that some important ABA-independent TFs involved in the specific plant response to abiotic stress combination have been identified. Considering that ABA levels dramatically change in response to environmental factors, the study of ABA-independent genes that are specifically regulated under stress combination may provide a marvelous tool for increasing plant resilience to climate change.

**SIGNIFICANCE STATEMENT:** This study in tomato Wt and ABA-deficient mutant plants reveals a specific and unique ABA-independent transcriptome reprogramming under abiotic stress combination, with the identification of some key TFs that were induced under these specific conditions. Taking into account that ABA levels dramatically change in all crops in response to environmental factors, the study of ABA-independent genes that are specifically regulated under stress combination may provide a marvelous tool for increasing plant resilience to climate change.

## INTRODUCTION

Changes to the earth’s climate, driven by increased anthropogenic emissions of heat-trapping greenhouse gases are already having widespread effects on the environment. The frequent and extreme weather conditions, such as hurricanes, heatwaves, wildfires, droughts, floods, and precipitation are dramatically increasing. These extreme events directly impact global agricultural production, resulting in losses of millions of dollars/euros every year (Mariani and Ferrante, 2017), crops with less average yields, and the unavoidable associated increase in food prices. The scientific community is aware of how abiotic stresses can dramatically affect plant growth and production, and for many decades, unceasing efforts have been made to deepen the identification on new molecular markers that can help plants to live and produce under a given abiotic stress. However, it has been vastly demonstrated that plants often have to face several abiotic stresses simultaneously (Mittler, 2006; Rivero *et al*., 2014; Zandalinas *et al*., 2021; Liu *et al*., 2023) and, under these conditions, the plant’s response significantly differs from that observed under single stresses. For instance, salinity applied in combination with heat induced a unique stress response in tomato plants that derived in a stronger plant protection, as compared to plants grown under salinity alone (Rivero *et al*., 2014). On the contrary, photosynthesis was more sensitive to the combination of soil waterlogging and high temperature than to the single stresses (Liu *et al*., 2023). The literature has also shown that elevated CO_2_ has the potential to mitigate the negative effects of water deficit stress on plants, indicating that these two factors act in unison to protect the plant from stress, rather than to exacerbate the situation (Shanker *et al*., 2022). On the other hand, in *Eucalyptus globulus* exposed to a combination of heat and drought stresses, a specific protective response was triggered, which was not activated when either drought or heat stress was applied individually (Correia *et al*., 2018). This specific response was extended at physiological, biochemical, and molecular levels (Rivero *et al*., 2014), also affecting the plant proteome, metabolome, and transcriptome reprogramming (Sharma *et al*., 2018).

Plants have evolved intricate signaling pathways that help them adapt to environmental changes, and one important regulator of plant responses to abiotic stresses is the phytohormone abscisic acid (ABA). Under drought, cold, or salt stress conditions, plants accumulate increased amounts of ABA, with drought stress having the most prominent effect. Thus, it is known that ABA plays important roles in the induction of plant tolerance to these stress conditions (Shinozaki and Yamaguchi-Shinozaki, 1997; Xiong *et al*., 2002; Finkelstein *et al*., 2002; Wang *et al*., 2018). However, although the ABA signaling system engages many stress-responsive genes, a large number of them do not respond to ABA treatments. This suggests the existence of ABA-independent stress-response pathways (Shinozaki and Yamaguchi-Shinozaki, 2000). During abiotic stress, such as drought, salinity, or high temperatures, ABA accumulates in various organs and tissues (Popova *et al*., 1995; Larkindale and Knight, 2002; Muhammad Aslam *et al*., 2022), and this ABA-dependent pathway has been the most studied, given its relationship with stress signaling mechanisms. The ABA-dependent pathway involves ABA binding to its receptor, PYR/PYL/RCAR, which triggers the release of PP2C phosphatases from the receptor complex (Santiago *et al*., 2009; Cutler *et al*., 2010). This, in turn, allows the activation of SnRK2 kinases, which phosphorylate downstream targets, including transcription factors such as ABF/AREB. These transcription factors then bind to ABA-responsive elements (ABREs) in target gene promoters, leading to changes in gene expression and physiological responses (Yoshida *et al*., 2019; Zhao *et al*., 2023). ABA-dependent responses include stomatal closure, inhibition of growth, and induction of stress-related genes. For example, ABA-induced stomatal closure is mediated by ABA binding to PYR/PYL/RCAR receptors in guard cells, leading to the activation of ion channels and pumps that result in a decrease in turgor pressure and ultimately stomatal closure (Fujii *et al*., 2007; Dittrich *et al*., 2019). On the other hand, the ABA-independent pathway involves signaling pathways that do not require ABA receptors or ABA signaling components. This ABA-independent pathway is still largely unknown due to the difficulty in studying it. TFs such as DREB2A/2B, AREB1, RD22BP1, and MYC/MYB, regulate ABA-responsive gene expression by interacting with their corresponding cis-acting elements DRE/CRT, ABRE, and MYCRS/MYBRS (Tuteja, 2007). CBF-type TFs are crucial in water and salt stress tolerance in higher plants, whereas it has been shown that DREB1 is stimulated by drought, salt stress, and exogenous ABA (Yang et al., 2012). Conversely, overexpressing DREB1 upregulated both ABA-independent and ABA-dependent stress-induced genes (*COR15a* and *rd29B*, respectively) in *Arabidopsis*, indicating an interaction between the ABA-dependent and -independent pathways (Yang *et al*., 2011). Another study carried out by Lee et al. (2017) showed that the *OsERF71* gene in rice was involved in an ABA-independent pathway, regulating drought resistance by controlling cell wall modifications. After *OsERF71* overexpression, roots modifications were sufficient for providing drought resistance phenotypes and increasing yield under drought stress, indicating that an ABA-independent signaling pathway induced drought tolerance mechanisms (Lee *et al*., 2017). In this line, *AtMYB60*, an R2R3-MYB gene expressed in guard cells, was negatively modulated during drought, and was involved in regulating stomatal movements. A mutant with an *AtMYB60* T-DNA insertion showed a reduction in stomatal opening, and its effects on water loss and transpiration rate during drought stress were observed (Cominelli *et al*., 2005). The regulation of this gene was described as ABA-independent regulation. Many R2R3-MYB TFs have been identified in *Glycine max* (He *et al*., 2020), *Arabidopsis* (Stracke *et al*., 2001), *Oryza sativa* (Katiyar *et al*., 2012), and *Solanum lycopersicum* (Zhao *et al*., 2014). Studies have successfully shown that they play important roles in the plant’s response to biotic and abiotic stresses (Abe *et al*., 2003; Denekamp and Smeekens, 2003; Nagaoka and Takano, 2003) and the regulation of secondary metabolite biosynthesis (Abe *et al*., 2003; Stracke *et al*., 2007), among many other developmental processes. Most of the studies conducted in these R2R3-MYB TF families have been categorized under the ABA-dependent pathway. In a study that attempted to identify and characterize the R2R3MYB family in tomato, by Zhao et al. (2014), the transcript abundances of the 51 *SlR2R3MYB* genes were investigated under NaCl (100 mM), low temperature (4 °C), and ABA (100 μM) treatments at the three-true-leaves stage, respectively (Zhao *et al*., 2014). Their results indicated that 19 genes (∼37.3%) responded to at least one treatment, which included 10 genes responding to the NaCl treatment, 9 genes to ABA, and 14 genes to low temperature. Among these genes, only 3 genes (*SlMYB62*, *SlMYB74* and *SlMYB102*) responded to all three treatments, and interestingly, 9 genes (*SlMYB2, SlMYB20, SlMYB21, SlMYB53, SlMYB78, SlMYB92, SlMYB108, SlMYB113,* and *SlMYB114)* did not respond to ABA. Also, some genes behaved in an opposite manner to their expression profile when subjected to different treatments. For example, *SlMYB64* was induced by high salinity, but was repressed by low temperature, and *SlMYB28* was induced by high salinity and ABA, respectively, but was repressed by low temperature (Zhao *et al*., 2014).

All the available literature regarding the identification of genes that take part in ABA-dependent and –independent pathways are based on studies conducted with a single stress condition. Therefore, and given that the plant’s responses to abiotic stress combinations are very specific and not able to be deduced, if they are not assayed and studied in detail, it is important to deepen the knowledge on the regulation of the plant’s transcriptome under abiotic stress combination, if we want to succeed in generating plants with increased resilience to climate change. Moreover, if we take into account that ABA levels dramatically change in response to the environment, the study of these ABA-independent genes can provide a marvelous tool to make advances on this objective.

In this study, tomato ABA-deficient mutants (*flacca, flc*), with a disrupted molybdenum cofactor (MoCo) sulfurase gene (Solyc07g066480), were treated with and without the exogenous application of ABA, and subjected to salinity, heat, or the combination of salinity and heat. The modification reaction that allows MoCo to be utilized for AO and XDH in the last step of ABA synthesis from abscisic aldehyde is catalyzed by MoCo sulfurase; thus *flc* tomato plants are Moco-sulfurase mutants with an impairment in all AO and XDH activities (Min et al., 2000; (Sagi *et al*., 1999; Sagi *et al*., 2002; Min *et al*., 2000), as they are ABA-deficient plants. It has been shown that NADH oxidase activity in tomato *flc* mutants provides molecular evidence that this activity is not dependent on the sulfuration state of MoCo. Therefore, an accumulation of ROS that was not dependent on the endogenous concentration of ABA was observed (Yesbergenova *et al*., 2005).

An in-depth RNA-seq analysis of our *flc* mutants revealed that the combination of salinity and heat induced a significant reprogramming of the tomato transcriptome, and also indicated that from the 685 genes specifically regulated under the combination of salinity and heat in our *flc* mutants, 463 genes followed an ABA-independent pathway regulation under these conditions. Among these genes, we identified 6 TFs belonging to the R2R3MYB family that were significantly upregulated. A protein-protein interaction network showed that *SlMYB50* and *SlMYB86* were directly involved in the upregulation of the flavonol biosynthetic pathway-related genes. These results confirm that the overexpression of *SlMYB50* and *SlMYB86* followed an ABA-independent pathway, driving the expression of some key flavonoid biosynthesis-related genes. More importantly, this regulation was specifically induced under the combination of salinity and heat in tomato plants.

## RESULTS AND DISCUSSION

### 1. ABA-deficient *flc* mutants showed a dramatic phenotype due to an impairment in stomatal regulation

ABA plays many functions in plants, including growth and development-related functions (Cutler *et al*., 2010). Thus, a basal amount of ABA is required for promoting plant development of tissues and organs. ABA is synthetized from a β-carotene as a derivate from an epoxy-carotenoid precursor, which, through a previous oxidative excision, produces xanthoxin (Parry *et al*., 1988). Xanthoxin is then converted to ABA by a number of ring modifications to generate abscisic aldehyde, which is then oxidized to ABA by the molybdenum-containing aldehyde oxidase (AO; EC 1.2.3.1) (Walker-Simmons *et al*., 1989; Leydecker *et al*., 1995). In this sense, several mutants with a reduced capacity to synthesize ABA through the mutation of the molybdenum cofactor (MoCo)-containing AO have been described, including *flc*. The *flc* mutation leads to complete loss of AO and xanthine dehydrogenase (XDH) activities in the shoot, while retaining minor but measurable activity in the roots, where detectable low levels of ABA accumulate (Sagi *et al*., 1999).

In our experiments, deficient-ABA tomato plants (*flc*) presented a very weak phenotype as compared to Wt under control conditions (Figure 1a), with 70% less biomass than the latter (Figure 1b), attributed to the lower endogenous ABA content in leaves (Imber and Tal, 1970). When salinity and heat were applied as combined stresses, both genotypes, Wt and *flç* experienced the highest biomass reduction, although it was noted that *flc* experienced a stronger growth inhibition (55%) as compared to Wt (40%) under abiotic stress combination (Figure 1b), indicating the important role of ABA in facing a combination of salinity and heat (Suzuki *et al*., 2016). Figure 1c shows a representative single stoma from both genotypes grown under control and under every stress condition applied (Figure S1). Wt showed a clear stomata closure under salinity or heat applied individually, although stress combination induced stomata opening (Figure 1c), which indicates the specificity of abiotic stress combination in stomatal regulation (Rivero *et al*., 2014; Suzuki *et al*., 2016; Zandalinas, Rivero, *et al*., 2016; Martinez *et al*., 2018). Stomatal conductance, however, was not affected in Wt plants under single or combined stress conditions as compared to Wt control (Figure 1d). On the other hand, *flc* mutants showed a general state of opened stomata under control or any stress condition applied (Figure 1c). As it has been extensively published, ABA plays an important role in stomata regulation (Zandalinas, Balfagón, *et al*., 2016; Yoshida *et al*., 2019; Hsu *et al*., 2021). Therefore, it was expected that the ABA deficiency imposed on *flc* mutants would lead to a poor control in stomata closure (Tal *et al*., 1970). The results showed that *flc* ABA-deficient tomato mutants displayed a marked tendency to wilt, due to excessive transpiration resulting from lack of control of stomatal closure. In this line, ABA concentration was 17 times lower in *flc* mutants as compared to Wt under any condition used (Figure 1e); it was noted, however, that ABA concentration, although at very low levels, was significantly higher in *flc* mutants grown under single or combined stress as compared to *flc* control plants (Figure 1e). The *flc* mutant is impaired in shoot AO activities, but retains detectable activities in the roots, where low levels of ABA were found, with this ABA potentially transported to the shoots and leaves (Sagi *et al*., 1999), which can explain the small amounts of this hormone found in our *flc* leaves. In parallel, stomatal conductance in *flc* mutants increased under single or combined stress, with the treatments of heat, and the combination of salinity and heat, inducing a higher stomatal conductance (Figure 1d). Bradford et al. (1983) observed almost twice the values of stomatal conductance in *flc* mutants as compared to Wt plants under control conditions, as also shown in our results (Bradford *et al*., 1983). However, this is the first time that stomatal conductance was studied in these *flc* mutants under abiotic stress combination, and our results showed a positive correlation between the increase observed in ABA concentration and the stomatal conductance (Figure 1d and e).

**Figure 1.**
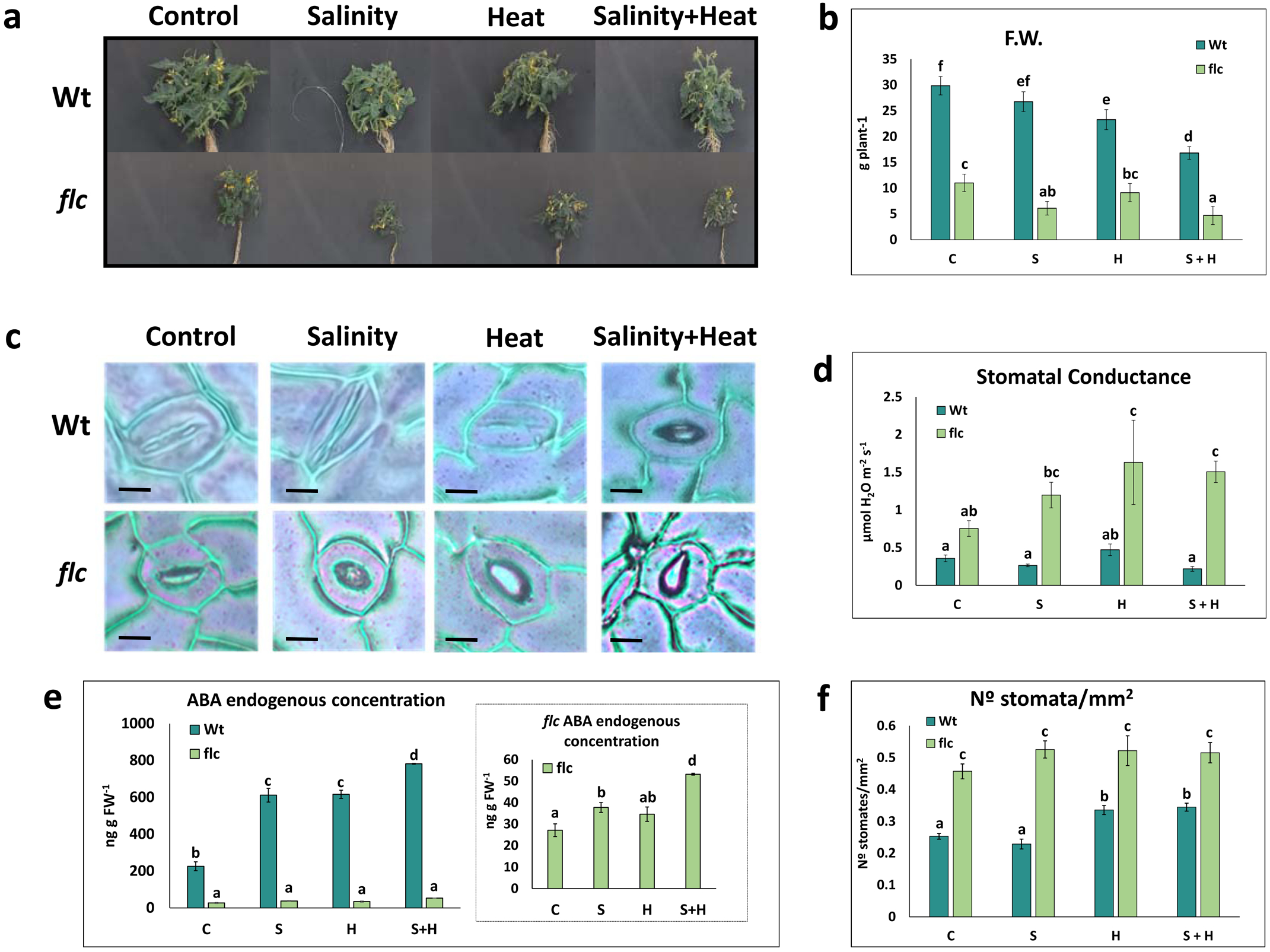
Physiological characterization in tomato wild-type (Wt) and *flacca* mutants (*flc*) plants under control (C), salinity (S, NaCl 100mM), heat (H, 35°C) and combination of salinity and heat (S+H). (a) Pictures of tomato plants at the sampling time (b) Whole plant fresh weight (FW) of tomato plants (c) Representative stoma of each of the genotype under the single or combined stress treatments. Whole micrographics for adaxial leaf surfaces can be found in Figure S1 (d) Stomatal conductance (e) ABA endogenous concentration (f) N° stomata per mm^2^. Values represent means ± SE (n = 6 in figure 1b and f, n=3 in figure 1d and e). Letters within each panel are significantly different at p < 0.05 according to Duncan’s test.

To verify if ABA was directly involved in the observed increase in stomatal conductance in *flc* mutants, or if it was due to an increase in the genotype-dependent stomata number, this parameter was measured (Figure 1d). While *flc* did not show any significant difference between the number of stomata counted under control or any stress condition used, Wt mutants showed a significant increase in the stomata number under heat, and the combination of salinity and heat, which was also in agreement with the increase in the stomatal conductance observed under these conditions (Figure 1d). Therefore, the small increase observed in ABA content in *flc* mutants grown under the combination of salinity and heat was perhaps not related with the significant increase observed in the stomatal conductance, as ABA levels were far below the standard limits, with this effect perhaps associated with the genotype-dependent increase in the stomata number per mm^2^, as compared to Wt, and the loss of stomata aperture control due to ABA deficiency.

### 2. ABA-deficient *flc mutants* showed a specific regulation under the combination of salinity and heat of 685 transcripts mostly related to cell catalytic, transcription regulation and transporter activities

To further investigate how ABA deficiency can affect the transcriptional machinery of the plants, an RNA-seq analysis was performed on Wt and *flc* grown under control, single, or combined stress conditions. Wt and *flc* mutants were grown under control conditions until all the plants had developed four true leaves, at which point the different stress treatments (single or combined) begun as described in the Material and Methods section. After 10 days under these conditions, metabolically-active and fully-expanded leaves were processed for RNA extraction, and RNA sequencing was performed by Macrogen Inc. (Seoul, Rep. of Korea) (Figure S2; Tables S1-S3). Wt control or *flc* control samples were used to normalize and obtain differentially expressed transcripts (DEGs) from Wt and *flc* plants, under single or combined stress (Table S4). Our results showed, considering all stress conditions, that ABA-deficient *flc* mutants had a lower total number of DEGs (5612), as compared to Wt (6423). This may indicate that *flc* mutants have a defective transcriptional machinery which might be affected by ABA-deficiency. It has been previously shown that ABA-dependent stress signaling first adjusts constitutively expressed TFs, leading to the expression of early response transcriptional activators, activating downstream stress tolerance effector genes (Hussain *et al*., 2021). This is consistent with the stronger growth inhibition observed in *flc* mutants under any stress condition applied, with it being more severe when salinity and heat were combined (Figure 1a and 1b). Also, it was clear from our results that the combination of salinity and heat induced a specific transcriptome response, in both Wt and *flc* mutants, which was completely different from that observed under salinity or heat applied individually, with 992 and 1243 specific DEGs, respectively (Figure 2a; Tables S5-S7). This observation has been extensively reported by other researchers, and highlights the importance of studying abiotic stress in combination (Rivero *et al*., 2014; Zandalinas, Rivero, *et al*., 2016; Lopez-Delacalle *et al*., 2021; Mikołajczak *et al*., 2022). However, from the specific DEGs found under the combination of salinity and heat, only 132 genes were common to Wt and *flc* mutants, which indicated a specific transcriptional reprogramming for ABA-deficient tomato plants, which still showed 1110 DEGs that were specifically regulated under these conditions (Figure 2a; Table S7).

**Figure 2.**
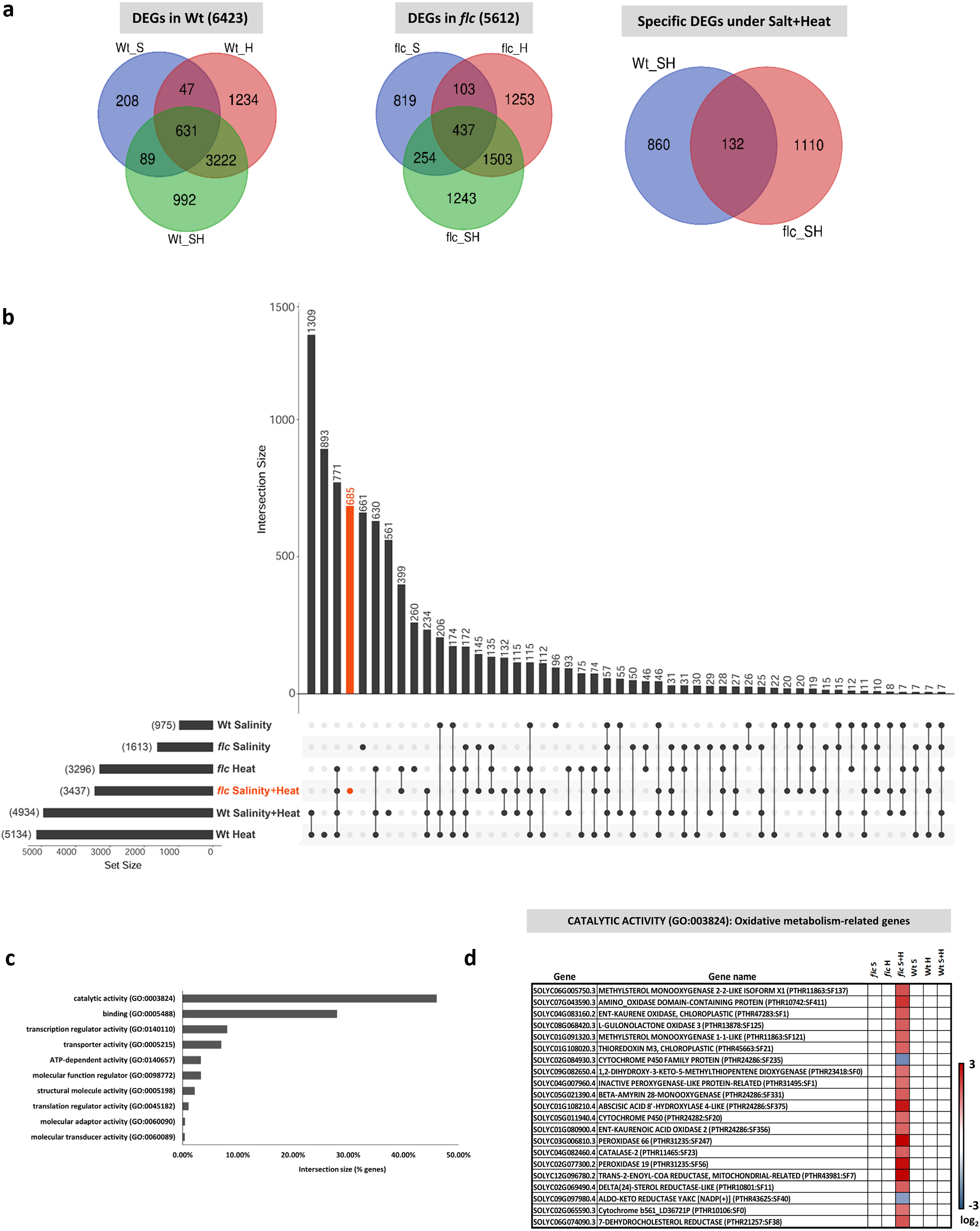
Differentially expressed genes (DEGs) specific to the combination of salinity+heat in *flacca* mutants. (a) Venn diagrams of the overlap between stress treatments in Wt and *flc* mutants and of the overlap between the specific DEGs found in Wt and *flc* under S+H (b) UpSet plot displaying the intersections between the sets of DEGs under single or combined stresses. The orange bar represents 685 DEGs specific of S+H in *flc* mutants (c) GO enrichment analysis of 685 DEGs specific of S+H combination in *flc* mutants (d) Heatmap of the specific oxidative metabolism-related genes found to be differentially expressed in *flc* mutants under S+H.

Our research interest was centered on the differentially expressed genes present in *flc* mutants under the combination of salinity and heat, and not under any other stress condition or in Wt. With this aim, DEGs obtained in both genotypes under single or combined stresses were used to create an UpSet plot (Figure 2b, Table S8). It was noted that the highest number of DEGs were obtained in Wt plants grown under heat or the combination of salinity and heat (5134 and 4934, respectively). However, only 1309 genes were shared by these two conditions (Wt-Heat and Wt-Salt+Heat), indicating the specificity of the stress combination on the development of the different plant response mechanisms. Also, Wt-Heat showed a large number of specifically induced DEGs (893), indicating the susceptibility of Wt plants to heat stress and the necessary plant reprogramming for inducing plant tolerance. It was noted that the Wt-Salinity treatment showed the lowest number of DEGs (975), with only 96 transcripts (∼10%) specific to this condition. This observation, together with the fact that Wt-Salinity shared only 22 transcripts with Wt-Heat, and only 55 transcripts with Wt-Salinity+Heat, indicate that Wt plants were more tolerant to salinity stress than to heat or the combination of stresses, and that plant’s metabolism need less readjustment under salinity stress alone (Rivero *et al*., 2014). This is an important finding to consider in the identification of specific molecular targets that can improve plant stress tolerance under field conditions, as it demonstrates that the single-stress studies may be incomplete and unsuccessful.

Following this line of study, the *flc* mutants showed a similar profile in the number of DEGs regulated under salinity, heat, or their combination, than those previously-described for Wt, with the highest number of DEGs found under heat or salinity+heat (3296 and 3437, respectively), and a lower number of DEGs under salinity alone (1613). Again, the reduced number of DEGs shared by *flc* mutants grown under heat and under salinity+heat (399) indicate the specificity of the stress combination on transcriptome reprogramming. When salinity was applied as an individual stress, 661 transcripts of the 1613 DEGs were specifically regulated by this condition in *flc* mutants (∼41%). This indicates that ABA-deficient *flc* mutants are more susceptible to salinity than Wt plants, and also, that ABA deficiency induced a salinity-specific salinity reprogramming that involved a higher number of differentially expressed transcripts.

Since the studies on abiotic stress combination can provide a better insight on specific molecular markers involved in plant stress tolerance under field conditions, we were particularly interested in the DEGs that were specifically regulated in ABA-deficient *flc* mutants grown under stress combination and, at the same time, that were not common with any other stress condition or with Wt plants. Thus, our next aim was to find new molecular markers that were associated with the specificity of the salinity and heat stress response, and whose regulation was independent of ABA, as there is a complete lack of research on this subject. The results from these studies can also facilitate the biotechnological applications of specific ABA-independent molecular markers for increasing plant resilience under environmental conditions where ABA can fluctuate. The UpSet plot showed a total of 685 DEGs that met these requirements (Figure 2b, marked in orange, Table S8). Interestingly, this specific condition provided the second highest number of DEGs (685), after Wt-Heat (893), that were not shared with any other genotype or condition used in our experiments, again indicating the specificity of this stress combination on ABA-deficient tomato plants suffering from this combined conditions. Suzuki et al. (2016) showed that *Arabidopsis* mutants deficient in abscisic acid metabolism and signaling were more susceptible to a combination of salt and heat stresses than wild type plants, which support our findings and demonstrates the significant plant transcriptional reprogramming under an ABA deficiency (Suzuki *et al*., 2016). We further extracted these 685 transcripts and a GO enrichment analysis was performed (http://www.pantherdb.org/geneListAnalysis.do), showing that most of them were related to catalytic (46%), transcription (40%), and transport activity (9%) (Figure 2c, Tables S9-S12). Among the DEGs with GO enrichment for catalytic activity, those related to oxidative metabolism were almost 20% of the total (Table S10), with most of them showing an upregulated expression under the combination of salinity and heat (Figure 2d, Table S13). It has been shown that ABA deficiency can trigger the expression of ROS-related genes to induce an ROS wave necessary for stomata control under abiotic stress in coordination with Ca^2+^ signaling waves (Mittler and Blumwald, 2015; Postiglione and Muday, 2020). An ABA deficiency in our *flc* mutants specifically induced the expression of some ROS-related transcripts under the combination of salinity+heat, but as these plants showed an absence in stomata control (Figure 1d and 1e), it might indicate that ABA levels are also fundamental for the Ca^2+^ waves and ROS signaling mechanisms to induce stomata closure (Mittler and Blumwald, 2015). DEGs related to binding and transcription regulation activity were also significantly enriched among the transcripts specifically regulated under the combination of salinity+heat (40%), confirming that ABA deficiency directly affects the transcriptional machinery reprogramming of tomato under the combination of salinity and heat.

Since ABA-deficient *flc* mutants showed a basal ABA concentration (very low, but with some ABA synthesis and accumulation, Figure 1e), it cannot be absolutely confirmed that the 685 DEGs found to be specifically regulated in *flc* under salinity and heat combination (Figure 2c), were also dependent on ABA-independent regulation. Therefore, we decided to perform a new experiment based on the exogenous supplementation of these *flc* mutants with 100 µM ABA (Figure 3), and to carry out a new RNA-seq analysis in these plants, to remove these ABA-dependent differentially regulated transcripts from the 685 DEGs found in *flc* mutants without ABA supplementation (Figure 3).

**Figure 3.**
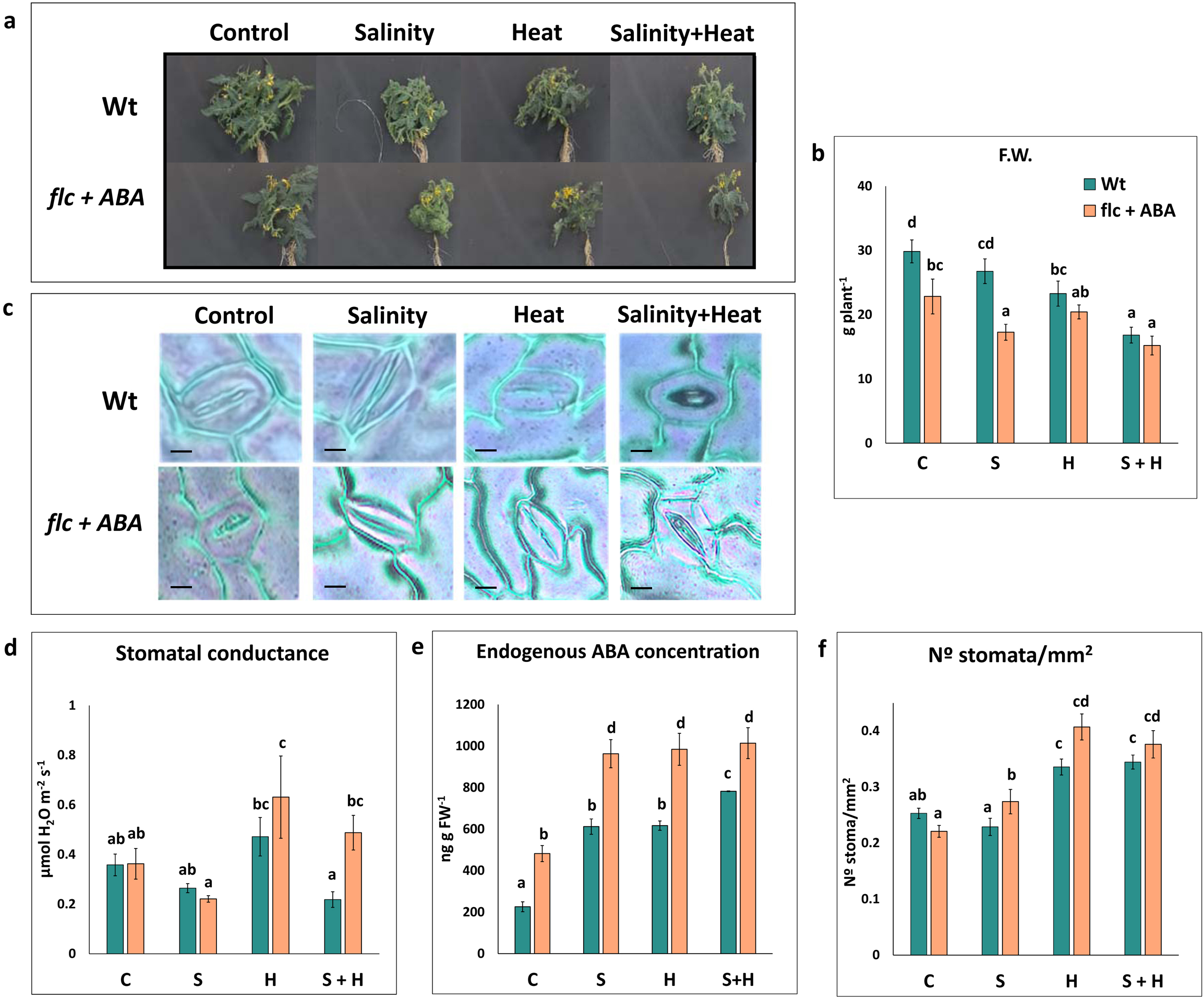
Physiological characterization of *flc* mutants with and without exogenous ABA application (100µM) under control, salinity (S, NaCl 100mM), heat (H, 35°C) and combination of salinity and heat (S+H). (a) Pictures of *flc* mutants at the sampling point (b) Whole plant fresh weight (FW) under single or combined stress with or without ABA supplementation (c) Representative stoma in *flc* mutants with or without ABA supplementation under single or combined stress. The whole micrograph pictures of adaxial leaf surface can be found in Figure S4 (d) Stomatal conductance in *flc* mutants with or without ABA supplementation under single or combined stress (e) ABA endogenous concentration in *flc* mutants with or without ABA supplementation under single or combined stress (f) N° stomata per mm^2^ measured within each condition used. Values represent means ± SE (n = 6 in figure 1b and f, n=3 in figure 1d and e). Letters within each panel are significantly different at p < 0.05 according to Duncan’s test.

### 3. Exogenous application of ABA to ABA-deficient *flc* mutant showed a Wt rescued phenotype under control or abiotic stress conditions

In this new experiment, we followed the same physiological/biochemical evaluation described in Figure 1. Thus, ABA-deficient *flc* mutants supplemented with 100 µM ABA were physiologically compared with Wt plants. For that, a preliminary experiment with different doses of ABA was previously performed with the *flc* mutants (Figure S3), with the dose of 100 µM exogenous ABA the one with the best fresh weight results. It was observed, however, that even with this exogenous application of ABA, *flc* mutants did not completely recover the Wt phenotype. Sagi et al. (1999) demonstrated that *flc* mutants had an impairment in NR activity as a consequence of a reduced water potential. This impaired the flow of phloem sap, which in turn compromised the level of total measured soluble sugar in *flc* roots. Under conditions of carbohydrate restriction, nitrate reduction is also reduced (Sagi *et al*., 1999). In any case, and since our aim was to revert the ABA content in *flc* mutants to Wt plants, the treatment of 100 µM of exogenous ABA was chosen for this (Figure S3). The different physiological measurements shown in Figure 1 were repeated here, obtaining very similar results between Wt and *flc* (Figure 3a-d), and confirming that the application of 100 µM ABA was successful. Also, the differences found in our previous experiment regarding biomass (Figure 1b), stomatal conductance (Figure 1d), and stomata number per mm^2^ (Figure 1f) were equalized by the exogenous application of ABA to our *flc* mutants (Figure 3b, 3d and 3f). In the same manner, the stomata in *flc* mutants were similar in morphology and aperture to those found in Wt under single or combined stress conditions (Figure 3c, Figure S4), although it was noted that stomatal conductance was only significantly higher in *flc* mutants grown under the combination of salinity and heat as compared to Wt (Figure 3d). Since stomata number per mm^2^ was, in this case, similar between Wt and *flc* mutants grown under any tested condition (except for the salinity treatment, which showed a slightly increase in the stomata number in *flc* mutants as compare to Wt) (Figure 3f), and the ABA endogenous concentration found in *flc* mutants was high enough to not be a limitation for stomata regulation (Figure 3e), the difference found in stomatal conductance under the combination of salinity and heat in Wt must be due to the specific environmental conditions (stress combination). Also, the endogenous concentration of ABA found in *flc* mutants (Figure 3e) ensured that any differentially regulated molecular markers found in these plants will include both ABA-dependent and ABA-independent genes.

Then, we performed a new RNA-seq analysis on the ABA-deficient *flc* mutants supplemented with 100 µM of ABA and grown under salinity, heat, or the combination of salinity and heat, following the same experimental period, sampling, total RNA extraction, and analysis protocols as those described in our previous experiment.

### 4. Specific ABA-independent transcriptome reprogramming occurring under the combination of salinity and heat induced the regulation of some transcription – related and serine/threonine kinase – related genes

The RNA-seq analysis performed in *flc* mutants supplemented with 100 µM ABA was performed in parallel with the previous RNA-seq analysis, and it is shown in Figure 4 and Tables S1-S3. After normalizing the data obtained in *flc*+ABA under single or combined stresses, against *flc+*ABA plants growing under control conditions, the DEGs obtained were matched with the 685 DEGs specifically expressed under the combination of salinity+heat found in our previous experiment (Figure 2c) using an UpSet plot (Figure 4a, Figure S14). As in our previous analysis, *flc*+ABA grown under salinity+heat or under heat alone obtained the highest number of DEGs (with 6583 and 5978 respectively). From these, 1326 transcripts were specifically regulated under the combination of salinity and heat, which again underlined the specificity of this stress combination on the plant’s transcriptional response.

**Figure 4.**
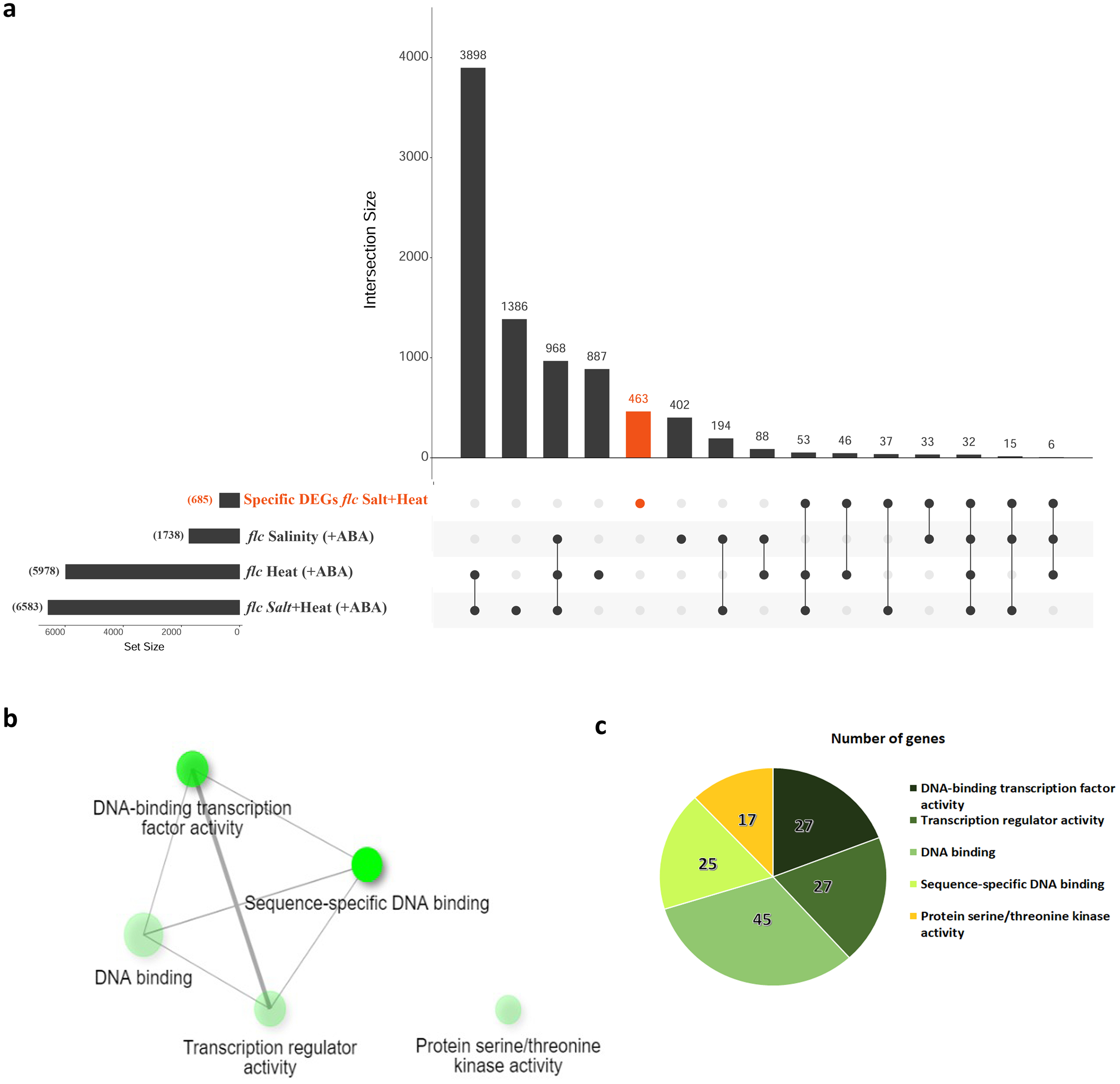
Identification of ABA-independent genes specifically regulated by the combination of salinity and heat. (a) UpSet plot displaying the intersections between the sets of specific DEGs found in *flc* mutants under the combination of salinity and heat (Figure 2) and DEGs found in *flc* mutants grown with ABA supplementation under single or combined stress. The orange bar represents 463 DEGs specific of the combination of salinity and heat and with an ABA-independent regulation (b) Molecular functions pathway enrichment of ABA-independent DEGs specific to the combination of salinity+heat (FDR cutoff: 0.02). The size of the circles is proportional to the number of genes related to that pathway and the color intensity is related to the pathway p-value significance level (p < 0.05) (c) Number of genes representative of each pathway. The raw data used for this figure can be found in Tables S14-S15.

By comparing the DEGs found in *flc* mutants+ABA under any of the stress conditions used, with the 685 DEGs shown in Figure 2c (*flc* mutants specifically regulated under the combination of salinity and heat), we can easily discard all the putative remaining ABA-dependent transcripts present in this last set of genes. Thus, a total of 222 transcripts were removed from the initial list of 685, and the remaining 463 genes were considered in this study to be specific to the combination of salinity and heat and, most importantly, with an ABA-independent regulation (Figure 4a). These 463 DEGs were subjected to an enriched network molecular function analysis (http://bioinformatics.sdstate.edu/go/; Figure 4b) using a FDR cutoff of 0.02, and a narrowed and enriched network containing a total of 124 transcription-related genes was found (Figure 4b-c, Table S15). Additionally, a group of 17 genes related to serine/threonine kinase activity proteins were also enriched among these 463 DEGs (Figure 4b-c, Table S15). Previously (Figure 2c) the GO enrichment analysis performed with the 685 DEGs that were specifically regulated in *flc* mutants grown under the combination of salinity+heat showed that a high percentage of these transcripts had a binding/transcription activity (∼40%), so it was not surprising that after removing these ABA-dependent DEGs from this first list, the remaining set of 124 genes appeared enriched in the new list. In the literature, we can find several publications that demonstrate the transcriptional regulation of gene expression governed by various key TF pathways that operate in ABA-dependent and -independent signaling pathways under abiotic stress (Hussain *et al*., 2021), which are regulated by AREB/ABFs (bZIP TFs) and DREB2A (AP2/ERF family of plant-specific TFs), respectively (Yoshida *et al*., 2014). DREB2 has been identified as the main TFs in ABA-independent pathways in *Arabidopsis* and soybean under drought and/or salinity stress applied individually (Yoshida *et al*., 2014; Chen *et al*., 2016). In addition, some studies have dealt with the functionality of these DREBs and the genes they may regulate under single abiotic stress conditions. However, as far as we know, this is the first evidence that shows an enriched list of transcription-related genes under ABA-independent regulation, which are specifically regulated under the combination of salinity and heat. And more importantly, these TFs were not significantly regulated under heat or salinity applied as individual stresses.

Of these 124 transcription-related genes, some of them were repeated among the different GO molecular functions groups, so only 49 true transcription-related transcripts were extracted and subjected to further analysis (Table S15).

### 5. *SlMYB50* and *SlMYB86* TFs were ABA-independently upregulated under the combination of salinity and heat and they were found to be related with flavonoid biosynthesis

Forty-nine transcription-related DEGs were found to be specifically regulated under the combination of salinity and heat, and they were also selected for being under ABA-independent regulation. These 49 transcription factors (TFs) were grouped under the different known TF families, and it was found that most of these families were represented, under our specific conditions, by 2-3 TFs, except for the MYB family, which were represented by 9 of them (Figure 5a, Table S16). Also, it was remarkable that most of the TFs found among all the TF families were upregulated, with only 4 TFs downregulated, indicating the need in *flc* mutants for transcriptional reprogramming within the ABA-independent pathway under the combination of salinity and heat. Members of the MYB family are involved in many cell events, including plant growth (Oppenheimer *et al*., 1991), plant responses to environmental factors and hormones (Urao *et al*., 1993; Magaraggia *et al*., 1997; Jin and Martin, 1999; Lee *et al*., 2007), signal transduction process (Gubler *et al*., 1995; Li *et al*., 2019; Hamaguchi *et al*., 2008), and pathogen defense (Shan *et al*., 2016; Noman *et al*., 2019), among others. It is also known that TFs from the MYB family may have an ABA-dependent or ABA-independent regulation (Zhao *et al*., 2014).

**Figure 5.**
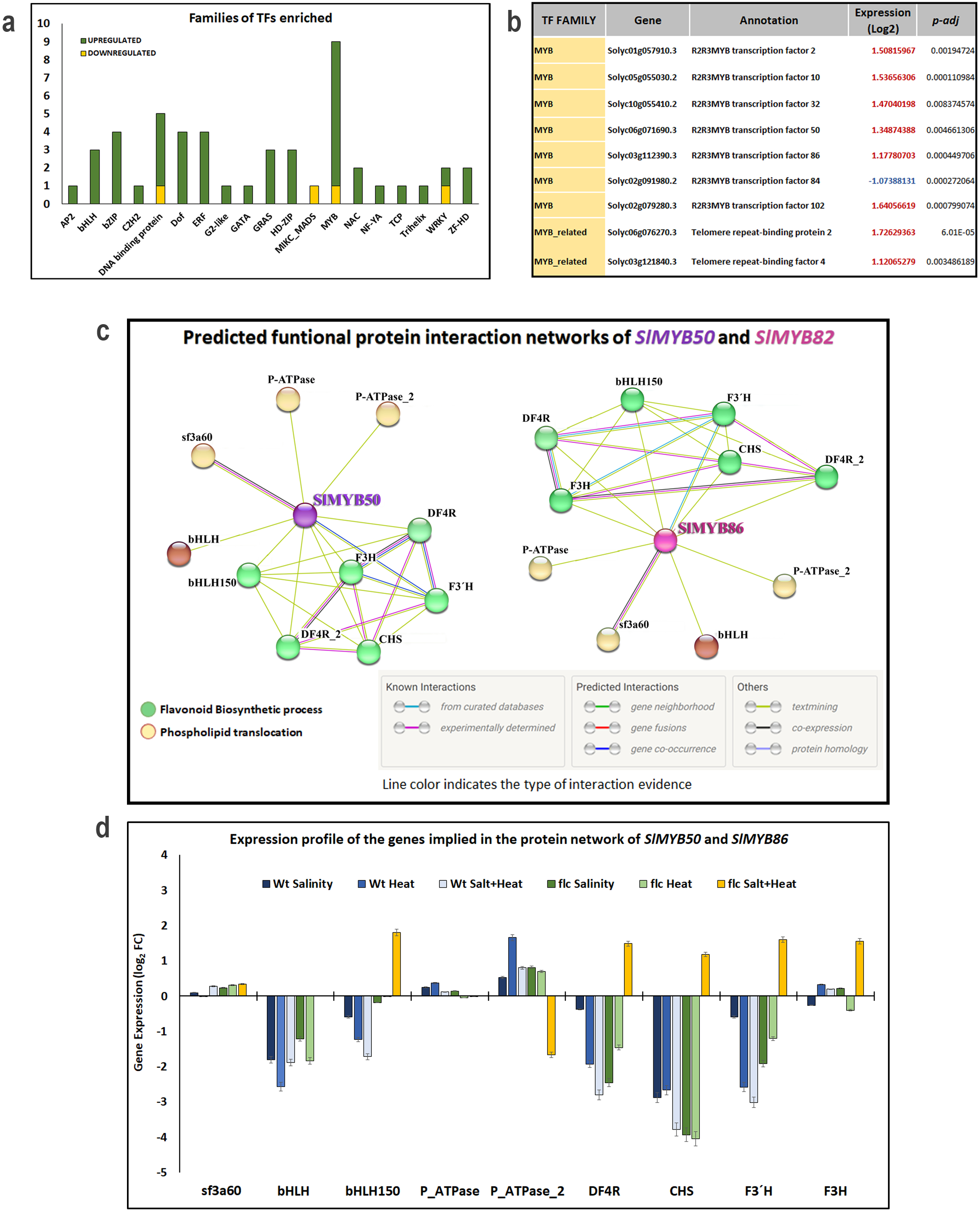
Identification of *SlMYB50* and *SlMYB86* transcription factors (TFs) with an ABA-independent regulation and specifically induced under the combination of salinity and heat. (a) Number of upregulated and downregulated transcription factors (TFs) identified for having an ABA-independent and specific regulation under the combination of S+H (b) MYB-related TFs identified in Figure 5a (c) Predicted functional protein interaction networks for SlMYB50 and SlMYB86 (d) Fold-change expression values (log_2_FC) of the genes represented within the SlMYB50 and SlMYB86 protein interaction networks in Wt and *flc* mutants under single and combined stresses. sf3a602: splicing factor subunit 3a, DF4R: dihydroflavonol 4-reductase, CHS: chalcone synthase, F3H: flavanone 3-hydroxylase and F3′H: flavonoid 3′-hydroxylase.

Our results showed that 6 genes containing the R2R3 MYB domain (*SlMYB2, SlMYB102, SlMYB10, SlMYB50, SlMYB32, SlMYB86*) were found to be upregulated, with the seventh (*SlMYB84)* repressed under our specific conditions (Figure 5b). R2R3MYB proteins comprise one of the largest families of transcription factors, and play regulatory roles in developmental processes and defense responses in plants (Zhao *et al*., 2014). A total of 121 R2R3MYB genes have been identified in the tomato genome. Tissue specificity or different expression levels of SlR2R3MYBs in different tissues suggest the differential regulation of tissue development, as well as metabolic regulation. Moreover, the transcript abundance level during abiotic conditions identified a group of R2R3MYB genes that responded to one or more stresses, suggesting that SlR2R3MYBs played major roles in the plant response to abiotic conditions, and were involved in signal transduction pathways (Zhao *et al*., 2014). It has been shown that the *Arabidopsis AtMYB2* was under ABA-dependent regulation after the application of drought and salinity stress as individual stresses (Abe *et al*., 2003). Also in rice, Yang et al. (2012) demonstrated that *OsMYB2*-overexpressing plants were more tolerant to salt, cold, and dehydration stresses, and more sensitive to abscisic acid than wild-type plants (Yang *et al*., 2012). However, our results indicate an ABA-independent regulation of this SlMYB2 under the combination of salinity and heat. On the other hand, Liao et al. (2008) showed that the transgenic *Arabidopsis* plants overexpressing soybean *GmMYB76, GmMYB92,* or *GmMYB177* showed a reduced sensitivity to ABA and exhibited better tolerance to salt and freezing treatments when compared with the Wt plants, following an ABA-independent regulation (Liao *et al*., 2008). In the same way *OsMYB3R2* was also found to be under ABA-independent regulation under cold, drought, and salinity stress (Dai *et al*., 2007). The results from that study suggested an ABA-independent regulation of some MYB R2R3 transcription factors under abiotic stress, but these experiments were assayed just under the application of a single stress and nothing was shown about their specific regulation when abiotic stresses were applied in combination.

To further delve into the putative role of the upregulation of these *SlMYB*-related transcription factors on the plant response to the combination of salinity and heat, a predicted functional protein interaction network analysis was performed using STRING (https://string-db.org/) with these 7 R2R3MYB genes (Figure S5). These predictive protein networks might help us to elucidate the potential targets of these TFs and their specific regulation. *SlMYB2* and *SlMYB102* were related to spliceosome; *SlMYB32* were related to biosynthesis of amino acids, and *SlMYB84* was related to DNA-binding function (Figure S9). Surprisingly, *SlMYB50* and *SlMYB86* showed an identical protein interaction network (Figure S6 and Tables S17 and S18) with a GO KEGG enrichment in the flavonoid synthesis pathway. The first hypothesis that emerged was that these two genes could probably be the same gene. However, after an identity check performed in the Solanaceae Genome Network database (https://solgenomics.net/tools/blast/) through a blast analysis (Figure S5), these two transcripts shared only 58.68% of identity (at the nucleotide and protein levels), and were therefore considered different and independent genes with an identical function. Thus, and due to the interest in studying the upregulation observed in these two transcription factors (*SlMYB50* and *SlMYB86*), which in turn may regulate the same genes, these two protein interaction networks were kept for further analyses. After performing a k-means clustering analysis within these two networks (Tables S17 and S18), and a GO enrichment analysis (Table S19), two specific pathways were found to be enriched in them: flavonoids biosynthetic process (green) and phospholipid translocation (yellow) (Figure 5c).

Flavonoids comprise a large group of secondary metabolites with a broad range of biological functions, including stress protection (Winkel-Shirley, 2002). The flavonoid biosynthesis pathway is a branch of the general phenylpropanoid pathway, with the first step catalyzed by chalcone synthase (CHS). In this pathway, the correlated activities of chalcone isomerase (CHI), flavanone 3-hydroxylase (F3H), flavonoid 3′-hydroxylase (F3′H), and flavonol synthase (FLS) result in the synthesis of the flavonols kaempferol and quercetin, which are further glycosylated by a number of glycosyltransferases (Winkel-Shirley, 2002). These flavonol glycosides are important products of the stress-activated flavonoid biosynthesis pathway, given their strong antioxidant activity. The regulation of the flavonoid biosynthesis pathway is largely at the level of transcription, both of the regulators and the biosynthetic genes (Quattrocchio *et al*., 2006; Jenkins, 2008). Thus, there is substantial interest on the underlying transcriptional regulatory networks that coordinate flavonoid biosynthesis (Broun, 2005; Quattrocchio *et al*., 2006; Jenkins, 2008), although very little or nothing is known about their pattern of synthesis and accumulation in plants that grow under abiotic stress combinations, or if this accumulation can follow an ABA-dependent and/or – independent regulation. A few years ago, our research group showed that most of the genes that belong to the flavonoid biosynthetic pathway were differentially and specifically upregulated under the combination of salinity and heat in tomato plants (Martinez *et al*., 2016). The fold-change expression values (log_2_) of the genes contained in these *SlMYB50* and *SlMYB86* protein interaction networks were measured in Wt and *flc* mutants under single and combined stresses (Figure 5d; Table S20). Surprisingly, those belonging to the flavonoids biosynthetic branch (DF4R, CHS, F3H and F3′H) were significantly upregulated in *flc* mutants grown under the combination of salinity and heat, whereas they were downregulated or not significant in Wt (under any stress condition applied), or in *flc* mutants grown under single stress conditions. These results confirm that the overexpression of *SlMYB50* and *SlMYB86* followed the ABA-independent regulation of these key flavonoids biosynthetic-related genes, and were specific to the combination of salinity and heat in tomato plants. Thus, *SlMYB50* and *SlMYB86* may be targeted as potential molecular markers for enhancing flavonoid biosynthesis and the plant’s resilience to the combination of salinity and heat in tomato plants. Moreover, *SlMYB50* and *SlMYB86* showed an ABA-independent regulation, and therefore, the biotechnological modification of their expression will perhaps not be affected by processes that rely on ABA-related signaling or, more importantly, by stresses that induce a major change in the cellular content of ABA.

## MATERIAL AND METHODS

### Plant growth and stress treatments

Wild-type (Wt) and the ABA-deficient mutants *flacca* (*flc*) (generously donated by TGRC, UC Davis, CA, USA, accession LA4479) of tomato (*Solanum lycopersicum* L. cv MicroTom) were used in all our experiments. *Flc* mutants are defective in the synthesis and/or maturation of the molybdenum-cofactor (MoCo) sulfurylase gene (Solyc07g066480), with these mutants deficient in ABA synthesis. Seeds were sterilized and germinated in vermiculite under optimal and controlled conditions (25/20°C day/night, light intensity of 400 μmol m^−2^ s^−1^, 16/8h day/night photoperiod, and 60-80% relative humidity). When the plants had at least two true leaves, 24 Wt plants and 48 *flc* mutants were transplanted into an aerated hydroponic system containing a modified Hoagland solution, and grown under controlled conditions until all of them developed four true leaves. The modified Hoagland solution had the following composition: KNO_3_ (6 mM), Ca(NO_3_)_2_ (4 mM), MgSO_4_ (1 mM), KH_2_PO_4_ (1 mM), KCl (50 μM), Fe-EDTA (20 μM), H_3_BO_3_ (25 μM), MnSO_4_⋅H_2_O (2 μM), ZnSO_4_⋅7H_2_O (2 μM), CuSO_4_⋅5H_2_O (0.5 μM), and (NH_4_)_6_Mo_7_O_24_⋅4H_2_O (0.5 μM) (Hoagland and Arnon, 1950). The pH of the nutrient solution was measured and maintained within 5.2–5.7, and the nutrient solution was renewed every three days.

When all the plants had developed four true leaves, half of the plants of each genotype (24 *flc* and 12 Wt) were transferred from Chamber A (where the plants were germinated and transplanted) to Chamber B, with the light, photoperiod and humidity parameters set identical to Chamber A, except for the environmental temperature variable, which was set to 35 °C, while the temperature remained at 25 °C in Chamber A. At the same time, half of the plants (12 *flc* and 12 Wt) of each genotype received 100 mM NaCl in their nutrient solution. Therefore, the experiment consisted in four treatments: control (25 °C and 0 mM NaCl), salinity (25 °C and 100 mM NaCl), heat (35 °C and 0 mM NaCl), and salinity + heat (35 °C and 100 mM NaCl). In parallel, half of our *flc* mutants of each treatment (24 plants, 6 plants per treatment) received an exogenous application of 100 µM ABA (based on a preliminary experiment, described below). After this time, all the plants were separated into roots and leaves, and fresh weight (FW) was recorded. The leaves of each plant were immediately stored at −80 °C for future determinations.

### ABA-exogenous application experimental design

Forty-eight *flc* mutant plants were germinated, transplanted, and grown under control conditions (25/20 °C day/night, light intensity of 400 μmol m^−2^ s^−1^, 16/8h day/night photoperiod, and 60-80% relative humidity), as described previously. When the plants had developed 4 true leaves, different concentrations of exogenous ABA (0, 50, 100, 150 µM) were applied to a group of 12 plants per ABA treatment. These treatments were repeated three times a week. Plants were kept under these conditions for 1 month, and were physiologically characterized (Figure S3). The application of 100 µM of exogenous ABA was chosen as the best for the stress combination experiments described previously.

### Stomatal conductance and stomata number determination

The stomatal conductance was measured every 3 days, as the stress treatments began in three single and fully-expanded leaves of each plant, using a LI-6400XT photosynthesis system (LICOR, Inc., Lincoln, NE, USA). Leaf gas exchange was measured in a 2 cm^2^ leaf cuvette. During these measurements, the conditions set in the LICOR were 1000 μmol photons m^−2^ s ^−1^, and 400 μmol mol^−1^ CO_2_. The leaf-air vapor pressure deficit was maintained between 1–1.3 kPa. The data reported are the mean 6 biological replicates per treatment and genotype.

Alternatively, the stomata number was determined by leaf epidermal impressions on 50 × 75 mm slides fashioned from 4.74 mm (3/16 in) cellulose acetate butyrate (CAB)/acetone. The impression of the adaxial (upper) part of the tomato leaf epidermis was performed as described by (Gitz and Baker, 2009) and photographed in a microscope (Olympus CKX41).

### ABA quantification

Endogenous ABA concentration was quantified in three biological samples per treatment and genotype in the Scientific and Technological Research Area (ACTI) of the University of Murcia. The extraction was carried out on 100 mg of frozen fresh leaf following the Müller and Munné-Bosch protocol (2011) with minor modifications (Müller and Munné-Bosch, 2011). Then, the organic extracts were injected into high-resolution UPLC-QToF-MS/MS equipment (Waters I-Class UPLC and Bruker Daltonics’ maXis impact Series QToF-MS with a resolution of ≥ 55,000 FWHM). Ionization by electrospray (ESI) was used as the source MS with both positive and negative polarities. Broad Band Collision Induced Dissociation (±bbCID) was used for MS/MS analysis with collision energies applied between 20 and 24 eV. Other parameters included a voltage source of 4.0 to 4.5 kV, a nebulizer pressure of 2.0 bar, drying temperature of 200 °C, a N_2_ flow rate of 9 L/min, and a mass range of 50-1200 Da. Sodium Formate (HCOONa) was used as a reference standard. An ABA standard curve (2-1000 ppm) was used for calculate the endogenous concentration of ABA in leaf samples.

### Total RNA extraction

Total RNA was isolated from 100 mg of frozen tomato leaves using the NucleoSpin® RNA kit (Reference no. 740949.50) in three biological replicates for each treatment and genotype. Total RNA was quantified and RNA integrity number (RIN) was checked in a bioanalyzer (2100 Expert Plant RNA Nano_DE13806178). Two µg of total RNA with a RIN ≥ 8 of each genotype and treatment were sent to Macrogen (Seoul) to proceed with the RNA-sequencing analysis.

### RNA sequencing, data analysis, functional annotation and GO enrichment analysis

RNA sequencing was performed in an Illumina HIseq (151 read-length paired-ends) analyzer. The raw reads obtained were filtered (Illumina passed-filter call) and further checked for sequence contaminants with fastQC (Table S1, S2, Figure S2). Contaminant-free, filtered reads for each sample were mapped with Bowtie/TopHat to the tomato genome sequence SL3.0 (ITAG4.0) (Hosmani *et al*., 2019) using HISAT2, which is known to handle spliced read mapping through the Bowtie2 aligner. After mapping, StringTie was used for transcript assembly. The expression profile was calculated for each sample and transcript/gene as read count, FPKM (Fragment per Kilobase of transcript per Million mapped reads) and TPM (Transcripts Per Kilobase Million) with a cutoff value of 0.1. DEG (Differentially Expressed Genes) analysis was performed on a comparison t-test pair using edgeR, where Wt and *flc* grown under the different stress treatments were compared with their respective control. The statistical analysis was performed using Fold Change, and exactTest per comparison pair. The significant results were selected on conditions of |fc|>=2 & exactTest raw p-value<0.05 (Table S3-S8). The DEG list was further analyzed with gProfiler (https://biit.cs.ut.ee/gprofiler/orth) and the Panther Classification System (http://www.pantherdb.org/geneListAnalysis.do;(Mi *et al*., 2019)) for gene set enrichment analysis (q-value 0.05) per biological process (BP), cellular component (CC), and molecular function (MF) (Table S9-S13). Upset plots were created in UpSetR (https://gehlenborglab.shinyapps.io/upsetr).

### Network enrichment analyses

Specific DEGs found in *flc* mutants under the combination of salinity and heat (Table S8) were submitted to the platform ShinyGO 0.77 (http://bioinformatics.sdstate.edu/go/ (Ge *et al*., 2020) with a FRD cutoff of 0.02 and a minimum pathway size of 5. The source code is available at https://github.com/iDEP-SDSU/idep/tree/master/shinyapps/go61.

For the construction of the protein-protein interaction network, the web-based tool STRING v11.5 (Szklarczyk *et al*., 2015), which is based on a large database of protein– protein interactions (PPI), was used. It also provides functionality for enrichment analysis of GO and protein domains in 5090 species (version 11), including tomato. The protein network analysis was performed based on the protein sequences as high confidence (0.7) with a maximum number of interaction of 10.

### Statistical analysis

All the experiments were performed in at least 3 biological repeats using the IBM SPSS 26 Statistics program. An analysis of variance (ANOVA) with a p-value < 0.05 was performed for fresh weight, stomatal conductance, stoma number per mm^2^, and ABA concentration, followed by a Duncan’s test. Standard error (SE) values for the different treatments and genotypes was calculated and added as shown in the figures. Significant changes in transcript expression as compared to the control (DEGs) were defined as log_2_ FC < -2 or >2 and adjusted P < 0.05 (negative binomial Wald test followed by Benjamini–Hochberg correction) (Ferreira and Zwinderman, 2006).

## SUPPLEMENTARY TABLES

**Table S1** Quatlity check of the reads obtained in the raw RNA-seq dataset for trimmed sequences.

**Table S2.** Mapping data statistics obtained in the RNA-seq analysis.

**Table S3.** Gene expression profile obtained for Wt and *flc* mutants. For the *flc* mutants, samples showed are without or with 100 µM ABA exogenous application. Reads are shown as counts, Fragments Per Kilobase Million (FPKM) and Transcripts Per Kilobase Million (TPM).

**Table S4.** Differentially expressed genes (DEGs) obtained in Wt and *flc* mutants under salinity, heat or the combination of salinity and heat. Samples were normalized against control condition.

**Table S5.** Differentially expressed genes for Wt showed in the Venn diagram of Figure 2a.

**Table S6.** Differentially expressed genes for *flc* mutants showed in the Venn diagram of Figure 2a.

**Table S7.** Comparison between the specific DEGs found under the combination of salinity and heat in Wt and *flc* mutants showed in Venn diagram for figure 2a.

**Table S8.** Gene IDs represented in the UpSet plot in Figure 2b for each comparison between Wt and *flc* mutants grown under single stress or stress combination.

**Table S9.** GO enrichment analysis of 830 DEGs from *flc* mutants specifically regulated under the combination of salinity and heat.

**Table S10.** GO enrichment analysis of DEGs with catalytic activity specifically regulated in *flc* mutants under the combination of salinity and heat.

**Table S11.** GO enrichment analysis of DEGs with catalytic activity specifically regulated in *flc* mutants under the combination of salinity and heat.

**Table S12.** GO enrichment analysis of DEGs with Transporter activity specifically regulated in *flc* mutants under the combination of salinity and heat.

**Table S13.** Oxidative metabolism-related genes contained in Table S10 (Catalytic activity). Expression data was taken from Table S4.

**Table S14.** Treatments comparison between DEGs found as specific in *flc* Salt + Heat (without ABA application) and DEGs in *flc* (with ABA application) under salinity, heat and Salt+Heat. Only 463 DEGs were found for not being shared with any *flc* stress treatment with ABA application (data represented in an UpSet plot in Figure 4a).

**Table S15.** Network enrichment analysis with the DEGs specifically regulated in *flc* mutants under the combination of salinity and heat and with an ABA-independent regulation.

**Table S16.** Transcription factor classification and fold-change found in *flc* mutants specifically regulated under the combination of salinity and heat and ABA-independent. **Table S17**. K-means clustering analysis of SlMYB50 protein interaction network (http://string-db.org).

**Table S18.** K-means clustering analysis of SlMYB86 protein interaction network (http://string-db.org).

**Table S19.** Process, Component, Clusters, KEGG, Compartments, Pfam and InterPro Enrichment in SlMYB50 and SLMYB86 protein interaction network analysis found in *flc* mutants.

**Table S20.** Gene expression of the genes contained in SlMYB50 and SlMYB86 protein interaction network. Data extracted from Table S4.

## SUPPLEMENTARY FIGURES

**Figure S1.** Light microscopy micrograph of adaxial leaf surface of tomato Wt and *flc* mutants grown under control, salinity, heat stress, or their combination. Scale bars: 100 μm.

**Figure S2.** Heatmap of the gene expression obtained for each sample and genotype in Wt and *flc* mutants grown under every stress condition applied. C: Control; S: Salinity; H: Heat; SH: Saliniity+Heat. 1, 2, and 3 represent the respective biological replication.

**Figure S3.** Phenotypes obtained for Wt and *flc* mutants grown under control conditions and with an exogenous application of ABA at different concentrations; (a) 0 μM ABA, (b) 50 μM ABA, (c) 100 μM ABA, and (d) 150 μM ABA

**Figure S4.** Figure S4. Light microscopy micrograph of adaxial leaf surface of tomato *flc* mutants grown with an exogenous ABA supplementation (100 μm) under control, salinity, heat stress, or their combination. Scale bars: 100 μm.

**Figure S5.** Protein interaction network for (1) *SlMYB2*, (2) *SlMYB102*, (3) *SlMYB10*, *SlMYB50*(4), (5) *SlMYB32*, (6) *SlMYB86*, (7) *SlMYB84*. The networks were constructed using STRING (https://stringldb.org/) for *Solanum lycopersicum* L., with a minimum interaction score of 0,7 (high confidence) and using UNIProt protein sequencing for each SlMYB transcription factor studied.

**Figure S6.** SlMYB50 and SLMYB86 identity check (https://solgenomics.net/tools/blast/) Proteins SlMYB50 and SLMYB86 showed a 58,68% of identity, therefore were considered as two different TFs with similar functions. They are able to regulate identical proteins in tomato metabolism, as shown in Figure 5C.

## ACKNOWLEDGMENTS

This research was supported by the Ministry of Economy and Competitiveness from Spain (Grant No. PGC2018-09573-B-100) to RMR; by the Ministry of Science and Innovation of Spain (Grant No. FPU20/03051) to MP-H, (Gran No. FPU21/01593) (Murcia, Spain) to SEM-L; and by University of Murcia Ph.D. contracts (Registry number 109144/2022) to JMM-G. We sincerely acknowledge Mario G. Fon for proof reading the manuscript. We thank ACTI (Scientific and Technological Research Area for University of Murcia) for the assistance with the analysis. All authors declare no commercial, industrial links or affiliations.

## AUTHOR CONTRIBUTIONS

MP-H did the experiments, data management, statistical analyses, analyzed RNA-seq data, wrote the manuscript and designed the figures and tables. SEM-L, JMM-G and IS contributed to the writing, editing, data analysis and literature updating. VA analyzed ABA endogenous concentration. RMR conceived the project, supervised, analyzed RNA-seq data, corrected and supervised with the writing and editing of the manuscript. All authors contributed to the article and approved the submitted version.

## DATA AVAILABILITY

The RNA-seq reads are available National Center for Biotechnology Information (NCBI) database under the Sequence Read Archive (SRA, https://submit.ncbi.nlm.nih.gov/subs/sra/), under the BioProject identification number PRJNA947059 (Submission ID: SUB12991832; direct link to datasets: https://www.ncbi.nlm.nih.gov/sra/PRJNA947059). Additionally, all the analyses are provided as row data as supplementary material (Figure S1-S5, Tables S1-S20).

## Notes

### Competing Interest Statement

The authors have declared no competing interest.

