## Supplementary material for "Specific ABA-independent tomato transcriptome reprogramming under abiotic stress combination": Figure S3

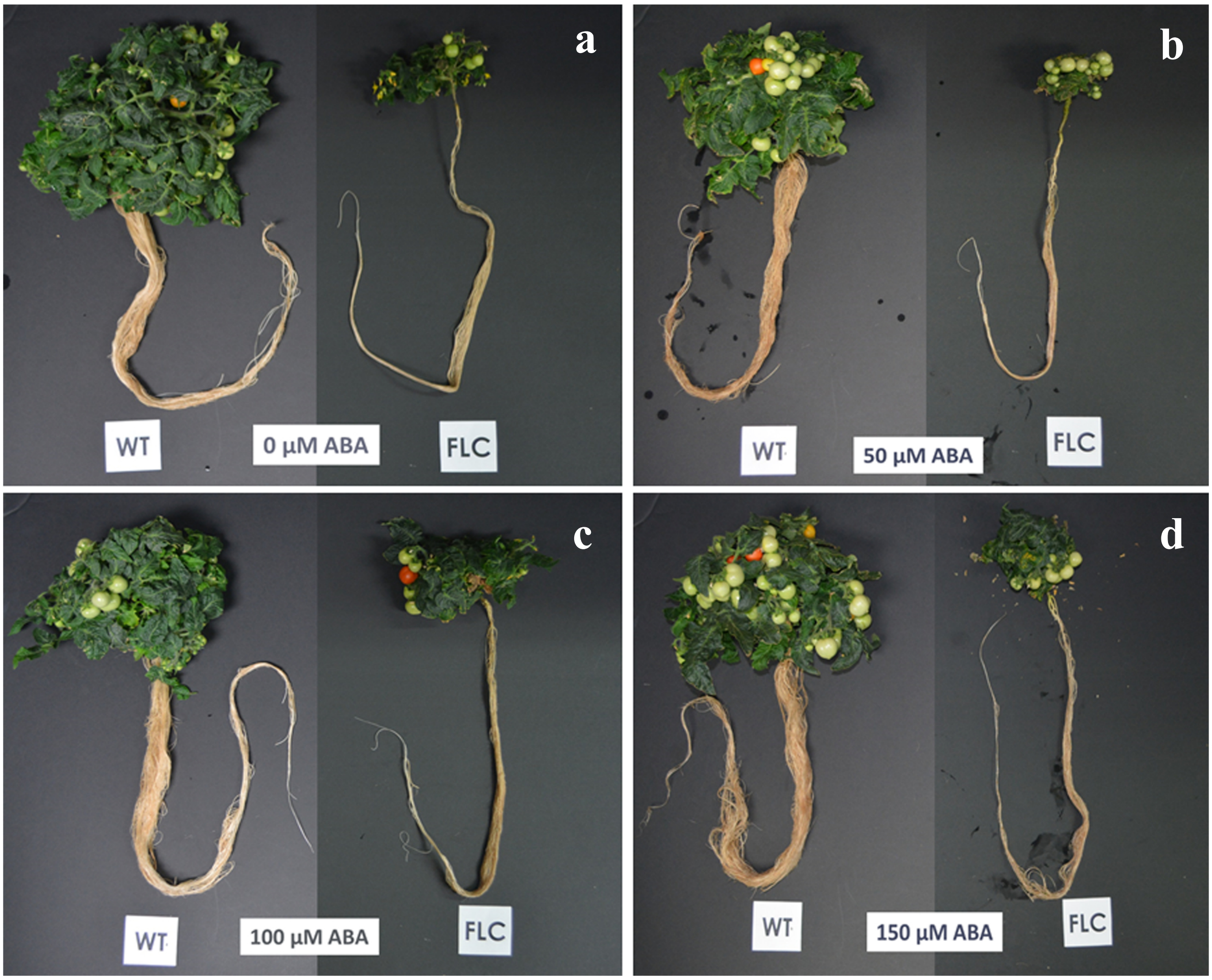


**Figure S3.** Phenotypes obtained for Wt and *flc* mutants grown under control conditions and with an exogenous application of ABA at different concentrations; (a) 0 µM ABA, (b) 50 µM ABA, (c) 100 µM ABA, and (d) 150 µM ABA
