## Supplementary material for "Specific ABA-independent tomato transcriptome reprogramming under abiotic stress combination": Figure S4

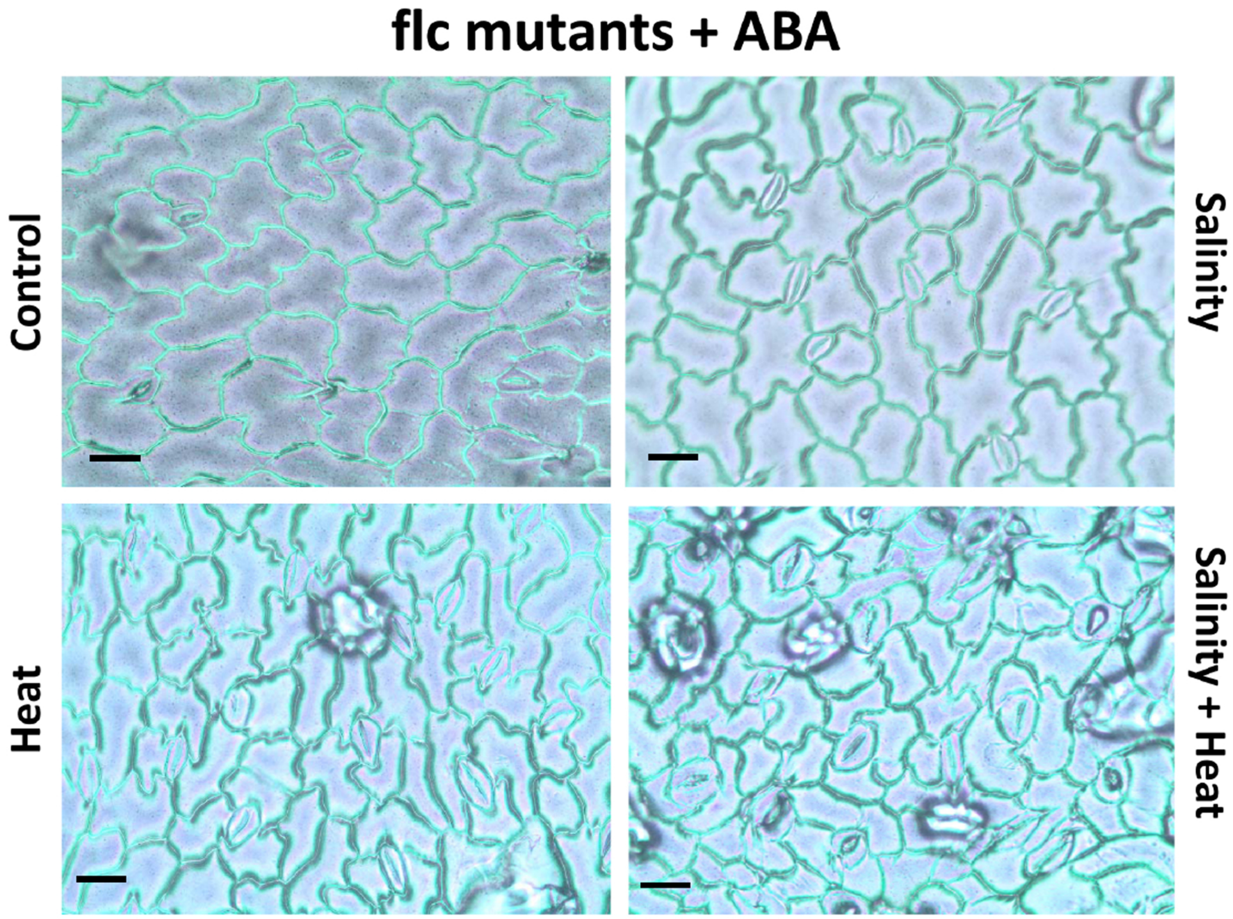


**Figure S4.** Light microscopy micrograph of adaxial leaf surface of tomato *flc* mutants grown with a exogenous ABA supplementation (100 µm) under control, salinity, heat stress, or their combination Scale bars: 100 µm
