## Supplementary material for "Specific ABA-independent tomato transcriptome reprogramming under abiotic stress combination": Figure S5

**Figure S5. Protein interaction network for (1) *SlMYB2*, (2) *SlMYB102*, (3) *SlMYB10*, *SlMYB50* (4), (5) *SlMYB32*, (6) *SlMYB86*, (7) *SlMYB84*.** The networks were constructed using STRING (https://string-db.org/) for *Solanum lycopersicum*, with a mínimum interaction score of 0,7 (high confidence) and using UNIProt protein sequencing for each *SlMYB* transcription factor studied

**1. SLMYB2**:

>Solyc01g057910.2.1
MDLRRGPWTVEEDLTLMNYIAHHGEGRWNTLARCAGLKRTGKSCRLRWLNYLRPDVRRGNITLEEQLLILELHSRWGN
RWSKIAQHLPGRTDNEIKNYWRTRVQKHAKQLKCDVNSKQFKDTMRYLWMPRLVERIQAAAGAGASTSSEVINPQNIV
QIDHNTCVTNSSTATASSDNSFGTPVSDLTDCCYNYPQQQQDCFQFVGESLISPTGYFHHALEFQGAAASAIDHQQNTTQ
WMDGANFSDNLWNIEDMWFLQQQLNNNVV

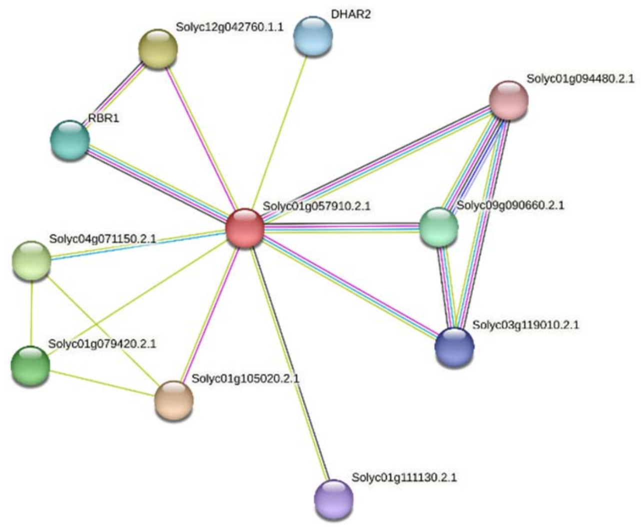


**Solyc12g042760. 1.1:** Leucine-rich repeat receptor-like protein kinase

**Solyc09g091280.1.2:** **RBR1** Retinoblastoma-related protein 1. GO:0008134 – transcription factor binding

**Solyc11g011250.1.1:** **DHAR2:** dehydroascorbate reductase 2. Glutathione dehydrogenase/transferase; Belongs to the GST superfamily.

**Solyc03g119010.2.1:** Pre-mrna-splicing factor spf27 homolog; Uncharacterized protein; Modifier of snc1,4

**Solyc01g111130.2.1:** Transcription factor bhlh121 isoform x2; Uncharacterized protein; Basic helix-loop-helix (bHLH) DNA-binding superfamily protein.

**Solyc01g105020.2.1:** Probable protein phosphatase 2c 62 isoform x1; Protein phosphatase 2C family protein.

**Solyc04g071150.2.1 :** Abscisic acid 8'-hydroxylase 3; Belongs to the cytochrome P450 family: enzyme encharged of the oxidative degradation of (+) ABA. Plays an important role in determining abscisic acid levels in dry seeds and in the control of postgermination growth.

**Solyc01g094480:** Pre-mRNA-splicing factor prp46 (AHRD V1 **-- B6K7I2_SCHJY); contains Interpro domain(s) IPR020472 G-protein beta WD-40 repeat, region

**Solyc09g090660:** Guanine nucleotide-binding protein subunit beta-like protein (AHRD V1 *--* GBLP_SCHPO); contains Interpro domain(s) IPR017986 WD-40 repeat, region

**Solyc01g079420.2:** Cytochrome c oxidase subunit VC family protein (AHRD V1 *-*- D7LT63_ARALY); contains Interpro domain(s) IPR008432 Cytochrome c oxidase subunit Vc

**2. SLMYB102**:

>Solyc02g079280.2.1
MGRAPCCDKNGLKKGPWTPEEDQKLIDYIQKHGYGNWRTLPKNAGLQRCGKSCRLRWTNYLRPDIKRGRFSF
EEEETIIQLHSILGNKWSAIAARLPGRTDNEIKNYWNTHIRKRLLRMGIDPVTHSPRLDLLDLSSILNHSIYNNSSH
HQMNLSRLLGHVQPLVNPELLRLATSLISSQRQNTNNFLIPNNLQENQIICQNQLPQMVQNNQIQDFSTISTTP
CVPFSSHEAQLMQPPITTKIEDFSSDLENFGNSQNNCQVINDDEWQLSNGVTDDYFPLQNYGYYDPLTSDENN
NNFNLQSVVLSNLSTPSSSPTPLNSNSTYFNNSSSTTTEDERDSYCSNMLNFDNIPNIWDTTNEFM*

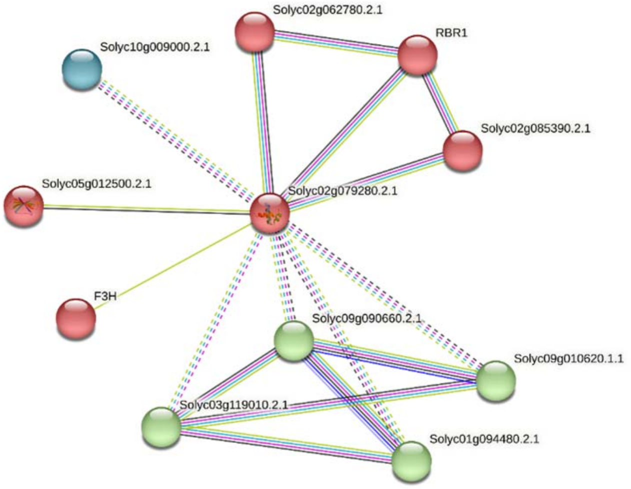


**Solyc02g079280.2.1:** R2R3MYB transcription factor 102
**Solyc10g009000.2.1:** Snrna-activating protein complex subunit isoform x1; snRNA activating complex family protein
**Solyc02g062780.2.1**: Chromodomain-helicase-DNA-binding protein 6
**Solyc09g091280.2.1:** Retinoblastoma-related protein 1 - RBR1
**Solyc02g085390.2.1:** Uncharacterized ATP-dependent helicase C25A8.01c
**Solyc09g010620.1.1:** Pre-mRNA-splicing factor
**Solyc09g090660.2.1:** Guanine nucleotide-binding protein subunit beta-like protein
**Solyc03g119010.2.1:** Pre-mrna-splicing factor spf27 homolog; Modifier of snc1,4
**Solyc09g090660.2.1:** Guanine nucleotide-binding protein subunit beta-like protein
**Solyc02g083860.2.1:** F3H - Naringenin 3-dioxygenase; Flavanone 3 beta-hydroxylase ; Belongs to the iron/ascorbate-dependent oxidoreductase family
**Solyc05g012500.2.1:** Probable wrky transcription factor 57; WRKY DNA-binding protein 57

### 3. SlMYB10:

>Solyc05g055030.1.1
MVRAPCCEKMGLKKGPWTQEEDQILINFIQKYGHENWRALPKQAGLLRCGKSCRLRWTNYLRPDIKRGNFSE
EEEQIIIKLHQLLGNRWSAIASRLPGRTDNEIKNFWHTHLKKRLEQSNNLASTITTKRRDEITTNASQNIDEHHIIS
SNYYQDSQNIVAYHPTYCHYQEDIQSNKEENMQYSTNYEHKDMTSFNNDMVFWYNVLMSSGNNASDEIS*

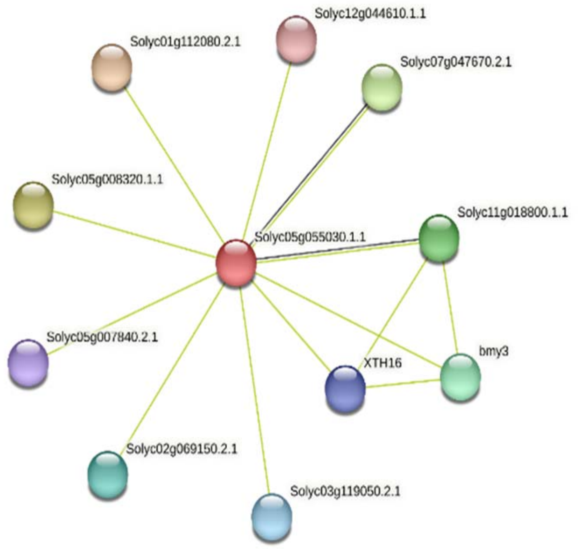


| | |
|---|---|
| **Solyc05g008320.1.1:** | Putative fasciclin-like arabinogalactan protein 20. |
| **Solyc01g112080.2.1:** | Lysm domain GPI-anchored protein 2 precursor |
| **Solyc12g044610.1.1:** | Transcription factor mybs3; Homeodomain-like superfamily protein. |
| **Solyc07g047670.2.1:** | Pescadillo homolog; Required for maturation of ribosomal RNAs and formation of the large ribosomal subunit. |
| **Solyc11g018800.1.1:** | Lignin-forming anionic peroxidase; Removal of H(2)O(2), oxidation of toxic reductants, biosynthesis and degradation of lignin, suberization, auxin catabolism, response to environmental stresses such as wounding, pathogen attack and oxidative stress. |
| **Solyc01g067660.2.1**, | bmy3: Beta-amylase; 1,4-alpha-glucan-maltohydrolase. |
| **Solyc07g052980.2.1**, | XTH16: Xyloglucan endotransglucosylase/hydrolase; Catalyzes xyloglucan endohydrolysis (XEH) and/or endotransglycosylation (XET). Cleaves and religates xyloglucan polymers, an essential constituent of the primary cell wall, and thereby participates in cell wall construction of growing tissues; Belongs to the glycosyl hydrolase 16 family |
| **Solyc03g119050.2.1**: | Myb domain protein 4r1 |
| **Solyc05g007840.2.1**: | Transcription termination factor mtef1, chloroplastic; Uncharacterized protein; Mitochondrial transcription termination factor family protein. |
| **Solyc02g069150.2.1:** | Vesicle-associated membrane protein 7B |

**4. SlMYB50**

>Solyc06g071690.2.1 Myb transcription factor (AHRD V1 *--* D6BV29_ORYSJ); contains Interpro domain(s) IPR015495 Myb transcription factor
MGRHSVFVKEKTRKGLWSPEEDEKLYNYITRFGVGCWSSVPKLAGLQRCGKSCRLRWINYLRPDLKRGMFSQEEEDMII
TLHKVLGNRWAQIAAKLPGRTDNEIKNFWNSNLKRKLIKQGIDPNTHKPLSENHQVRNEPNCTDKTSSLLMPKLPNMS
DSAEIQQPFHFFNSKRNFNSQAVTRELTEVSKNQLVSKQVFDPLFLYEFQASVNPIGPYAHHHNQIEGNQDFGFCSNFQ
HVHMTTESDISDSSTSRMSTSNSSNTMISHYNSVGIQMNEMLEWDADNKIDSLIQYPYVGIKNEENYSNNNNTLSGEN
LDVFHHI*

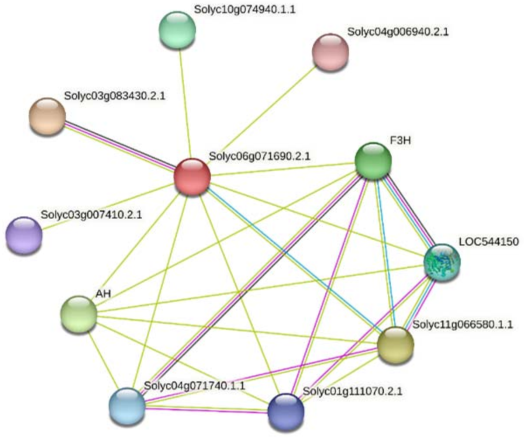


| | |
|---|---|
| **Solyc03g007410.2.1:** | Transcription factor speechless; Uncharacterized protein; Basic helix-loop-helix (bHLH) DNA-binding superfamily protein. |
| **Solyc03g083430.2.1:** | Splicing factor sf3a60 homolog; Uncharacterized protein; Splicing factor-related. |
| **Solyc10g074940.1.1:** | Phospholipid-transporting atpase 1-like; Belongs to the cation transport ATPase (P-type) (TC 3.A.3) family. Type IV subfamily. |
| **Solyc04g006940.2.1:** | Putative phospholipid-transporting atpase 9; Belongs to the cation transport ATPase (P-type) (TC 3.A.3) family. Type IV subfamily. |
| **Solyc02g085020.2.1:** | **LOC544150**: **(DFRA)** Bifunctional dihydroflavonol 4-reductase/flavanone 4-reductase; Dihydroflavonol 4-reductase; Bifunctional enzyme involved in flavonoid metabolism. |
| **Solyc11g066580.1.1:** | **(F3´H)** Flavonoid 3`,5`-hydroxylase; Putative flavonoid 3'5' hydroxylase ; Belongs to the cytochrome P450 family. |
| **Solyc01g111070.2.1:** | **(CHS)** Type iii polyketide synthase b; Chalcone and stilbene synthase family protein; Belongs to the chalcone/stilbene synthases family. |
| **Solyc04g071740.1.1:** | **(DFRA)** Dihydroflavonol 4-reductase. DF4R |
| **Solyc09g065100.1.1:** | **(AH):** Basic helix-loop-helix protein a isoform x1; Basic helix-loop-helix (bHLH) DNA-binding superfamily protein. |
| **Solyc02g083860.2.1:** | **(F3H):** Naringenin 3-dioxygenase; Uncharacterized protein; Flavanone 3 beta-hydroxylase ; Belongs to the iron/ascorbate-dependent oxidoreductase family. |

**5. SlMYB32**:

>Solyc10g055410.1.1

(AHRD V1 **-- B9H191_POPTR); contains Interpro domain(s) IPR017930 Myb-type HTH DNA-binding domain

MGRSPCCEKAHTNKGAWTKEEDERLISYIRAHGEGCWRSLPKAAGLLRCGKSCRLRWINYLRPDLKRGNFTEEEDELII
KLHSLLGNKWSLIAGRLPGRTDNEIKNYWNTHIRRKLLSRGIDPTTHRSINDPTTIPKVTTITFAAAHENIKDIDQQDEMI
NIKAEFVETSKESDNNEIIQEKSSSCLPDLNLELRISPPHHQQLDHHRHHQRSSSLCFTCSLGIQNSKDCSCGSESNGNG
WSNNMVSMNIMAGYDFLGLKTNGLLDYRTLETK*

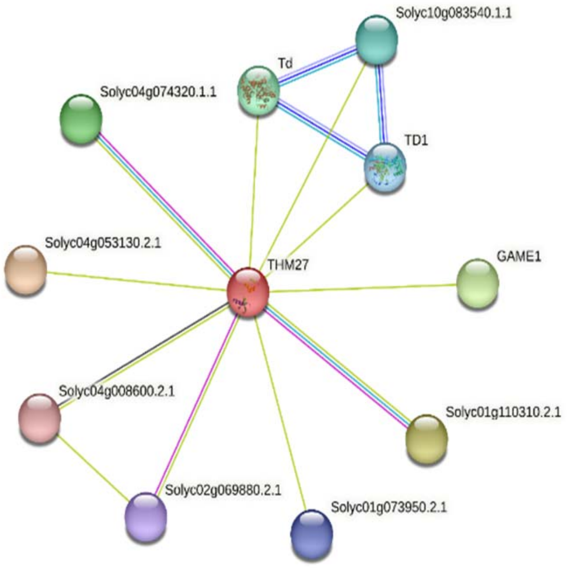


| | |
|---|---|
| **Solyc09g008670.2.1:** | **Td:** Threonine dehydratase biosynthetic, chloroplastic ; Belongs to the serine/threonine dehydratase family. |
| **Solyc10g083540.1.1:** | Threonine dehydratase I |
| **Solyc10g083760.1.1:** | **TD1:** Chloroplast threonine deaminase 1 precursor; Has a housekeeping role in isoleucine biosynthesis (Probable) |
| **Solyc07g043490.1.1:** | **GAME1:** Udp-galactosyltransferase precursor; Glycosyltransferase; Glycoalkaloid metabolism 1 ; Belongs to the UDP-glycosyltransferase family |
| **Solyc01g110310.2.1:** | GATA transcription factor; Transcriptional activator that specifically binds 5'- GATA-3' or 5'-GAT-3' motifs within gene promoters |
| **Solyc01g073950.2.1:** | Bromodomain protein |
| **Solyc02g069880.2.1:** | E3 ubiquitin-protein ligase DIS1 isoform X1; E3 ubiquitin-protein ligase; E3 ubiquitin-protein ligase that mediates ubiquitination and subsequent proteasomal degradation of target proteins. E3 ubiquitin ligases accept ubiquitin from an E2 ubiquitin- conjugating enzyme in the form of a thioester and then directly transfers the ubiquitin to targeted substrates. |
| **Solyc04g008600.2.1:** | Chromosome segregation in meiosis protein 3; Plays an important role in the control of DNA replication and the maintenance of replication fork stability |
| **Solyc04g053130.2.1:** | Stress enhanced protein 2, chloroplastic; Stress enhanced protein 2 |
| **Solyc04g074320.1.1:** | Protein transparent testa 1; C2H2 and C2HC zinc fingers superfamily protein |

### 6. SlMYB86:

>Solyc03g112390.2.1 Myb transcription factor (AHRD V1 *--- D6BV29_ORYSJ); contains Interpro domain(s) IPR015495 Myb transcription factor
MGRHSCSVKQKLRKGLWSPEEDEKLSNYITNFGIGSWSSVPKLAGLQRCGKSCRLRWINYLRPDLKRGMFSQDEE
DKIISLHQVLGNRWAQIAAQLPGRTDNEIKNFWNSSLKKKLMKQGIDPNTHKPLKENQIIKDEENCTNKTSMLQIP
PHLNEMGNGQFTDSKQVFDLLFVHDFQSNTNPREYNSQVLAQYHDHQGEFENHQNYVFCSGSSVTKLEHVQMT
ETDFGSSSTSRMSSSNSSNMCSNQNTAGIQINGMSENSEALSWDIENKMESLFQYPYIGIKNEESKSSPSQERDQL
YGNTTSGDFMSNYPLSSLTEEFKWG*

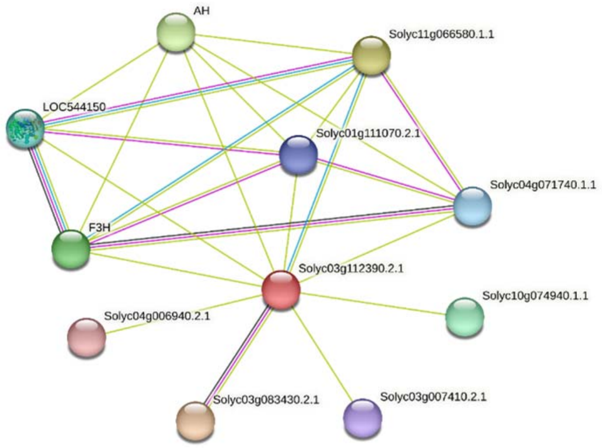


| | |
|---|---|
| **Solyc03g007410.2.1:** | Transcription factor speechless; Uncharacterized protein; Basic helix-loop-helix (bHLH) DNA-binding superfamily protein. |
| **Solyc03g083430.2.1:** | Splicing factor sf3a60 homolog; Uncharacterized protein; Splicing factor-related. |
| **Solyc10g074940.1.1:** | Phospholipid-transporting atpase 1-like; Belongs to the cation transport ATPase (P-type) (TC 3.A.3) family. Type IV subfamily. |
| **Solyc04g006940.2.1:** | Putative phospholipid-transporting atpase 9; Belongs to the cation transport ATPase (P-type) (TC 3.A.3) family. Type IV subfamily. |
| **Solyc02g085020.2.1:** | **LOC544150**: Bifunctional dihydroflavonol 4-reductase/flavanone 4-reductase; Dihydroflavonol 4-reductase; Bifunctional enzyme involved in flavonoid metabolism. |
| **Solyc11g066580.1.1:** | Flavonoid 3`,5`-hydroxylase; Putative flavonoid 3'5' hydroxylase ; Belongs to the cytochrome P450 family. |
| **Solyc01g111070.2.1:** | Type iii polyketide synthase b; Chalcone and stilbene synthase family protein; Belongs to the chalcone/stilbene synthases family. |
| **Solyc04g071740.1.1:** | Dihydroflavonol 4-reductase. DF4R |
| **Solyc09g065100.1.1:** | **AH:** Basic helix-loop-helix protein a isoform x1; Basic helix-loop-helix (bHLH) DNA-binding superfamily protein. |
| **Solyc02g083860.2.1:** | **F3H:** Naringenin 3-dioxygenase; Uncharacterized protein; Flavanone 3 beta-hydroxylase ; Belongs to the iron/ascorbate-dependent oxidoreductase family. |

**7. SlMYB84**

>Solyc02g091980.1.1

MGRAPCCDKANVKRGPWSPEEDSKLKAYIEQHGTGGNWITLPQKVGLKRCGKSCRLRWLNYLRPNIKHGEFTDE
EDNIICTLYMSIGSRWSIIAAQLPGRTDNDIKNYWNTRLKKKLLLCNKQRKDQRPRTGSYINHNKLEMMNEHENFF
ATHQTINNAYSWPSQQILFSTLIAPQNHDLAESSSANSHNFQYSSTDQVYFSQDQLCQISSTNQLPSMNLVNGNTC
NVISNGYYPSNGVVINNGLQEYNNYNSVGLGHDALNSTSTTHSQQVDKSVLEMVNSSTISTTSQDQSTSWEELSPL
VNYPPSVFETVSPYYVFEEQRYMGLLKQ*

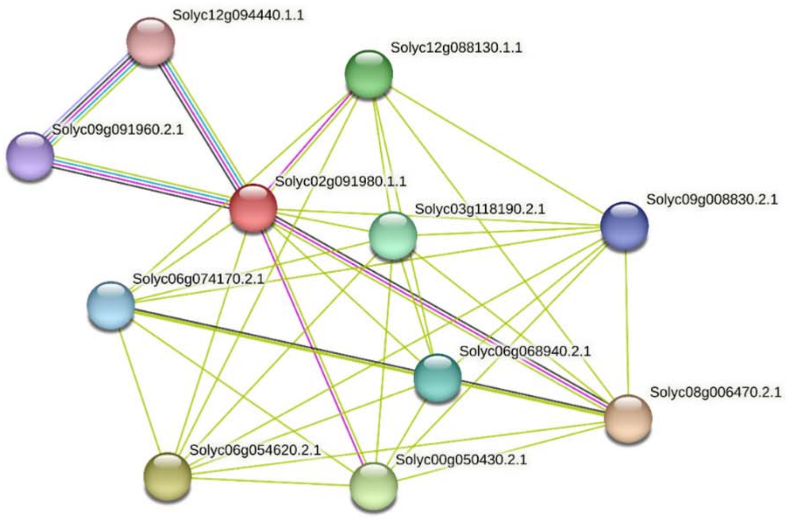


**Solyc02g091980.1.1:** R2R3MYB transcription factor 84
**Solyc09g091960.2.1:** HMG (high mobility group) box protein with ARID/BRIGHT DNA-binding domain
**Solyc12g094440.1.1:** HMG (high mobility group) box protein with ARID/BRIGHT DNA-binding domain
**Solyc12g088130.1.1:** BHLH transcription factor
**Solyc03g118190.2.1**: Ethylene responsive transcription factor 2a
**Solyc09g008830.2.1:** Os01g0318400 protein (Fragment) (AHRD V1 *-*- Q7XXQ7_ORYSJ)
**Solyc06g068940.2.1:** C2 domain-containing protein
**Solyc08g006470.2.1:** Zinc finger family protein C2H2-type
**Solyc00g050430.2.1:** bHLH transcription factor 073
**Solyc06g054620.2.1:** Zinc finger CCCH domain-containing protein 34
**Solyc06g074170.2.1:** NAC domain protein IPR003441
