## Supplementary material for "Specific ABA-independent tomato transcriptome reprogramming under abiotic stress combination": Figure S6

**Supplemental Figure S6. SlMYB50 and SLMYB86 identity check (https://solgenomics.net/tools/blast/)**

Proteins SlMYB50 and SLMYB86 showed a 58,68% of identity, therefore were considered as two different TFs with similar functions. They are able to regulate identical proteins in tomato metabolis, as shown in Figure 5C.

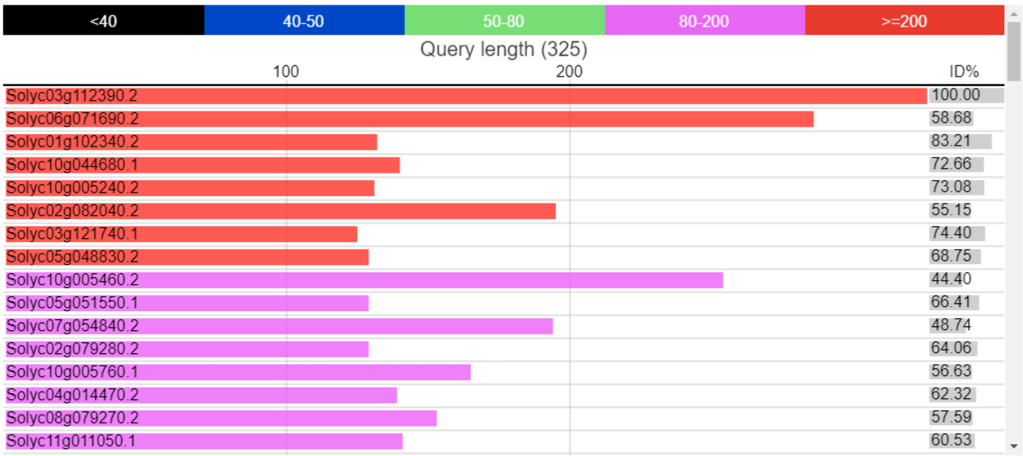


Solyc03g112390.2.1 vs Tomato Genome protein sequences (ITAG release 2.40)

| SubjectId | id% | Aln | evalue | Score | Description |
|---|---|---|---|---|---|
| Solyc03g112390.2 | 100.00 | 325/325 | 0.0 | 678 | R2R3MYB transcription factor 86 SlMYB86Length=325 |
| Solyc06g071690.2 | 58.68 | 186/317 | 1e-123 | 357 | R2R3MYB transcription factor 50 SlMYB50Length=321 |

```
>Solyc06g071690.2 R2R3MYB transcription factor 50 SlMYB50
Length=321

 Score = 357 bits (916),  Expect = 1e-123, Method: Compositional matrix adjust.
 Identities = 186/317 (59%), Positives = 221/317 (70%), Gaps = 51/317 (16%)

Query  1    MGRHSCSVKQKLRKGLWSPEEDEKLSNYITNFGIGSWSSVPKLAGLQRCGKSCRLRWINY  60
            MGRHS  VK+K RKGLWSPEEDEKL NYIT FG+G WSSVPKLAGLQRCGKSCRLRWINY
Sbjct  1    MGRHSVFVKEKTRKGLWSPEEDEKLYNYITRFGVGCWSSVPKLAGLQRCGKSCRLRWINY  60

Query  61   LRPDLKRGMFSQDEEDKIISLHQVLGNRWAQIAAQLPGRTDNEIKNFWNSSLKKKLMKQG  120
            LRPDLKRGMFSQ+EED II+LH+VLGNRWAQIAA+LPGRTDNEIKNFWNS+LK+KL+KQG
Sbjct  61   LRPDLKRGMFSQEEEDMIITLHKVLGNRWAQIAAKLPGRTDNEIKNFWNSNLKRKLIKQG  120

Query  121  IDPNTHKPLKENQIIKDEENCTNKTSMLQIPP----------------------------  152
            IDPNTHKPL EN  +++E NCT+KTS L +P
Sbjct  121  IDPNTHKPLSENHQVRNEPNCTDKTSSLLMPKLPNMSDSAEIQQPFHFFNSKRNFNSQAV  180

Query  153  --HLNEMGNGQFTDSKQVFDLLFVHDFQSNTNPREYNSQVLAQYHDHQGEFENHQNYVFC  210
               L E+   Q   SKQVFD LF+++FQ++ NP       +  Y  H  + E +Q++ FC
Sbjct  181  TRELTEVSKNQLV-SKQVFDPLFLYEFQASVNP-------IGPYAHHHNQIEGNQDFGFC  232

Query  211  SGSSVTKLEHVQM-TETDFGSSSTSRMSSSNSSN-MCSNQNTAGIQINGMSENSEALSWD  268
            S       +HV M TE+D   SSTSRMS+SNSSN M S+ N+ GIQ+N      E L WD
Sbjct  233  SN-----FQHVHMTTESDISDSSTSRMSTSNSSNTMISHYNSVGIQMN------EMLEWD  281

Query  269  IENKMESLFQYPYIGIK  285
             +NK++SL QYPY+GIK
Sbjct  282  ADNKIDSLIQYPYVGIK  298
```
