## Supplementary material for "Specific ABA-independent tomato transcriptome reprogramming under abiotic stress combination": Figure S1

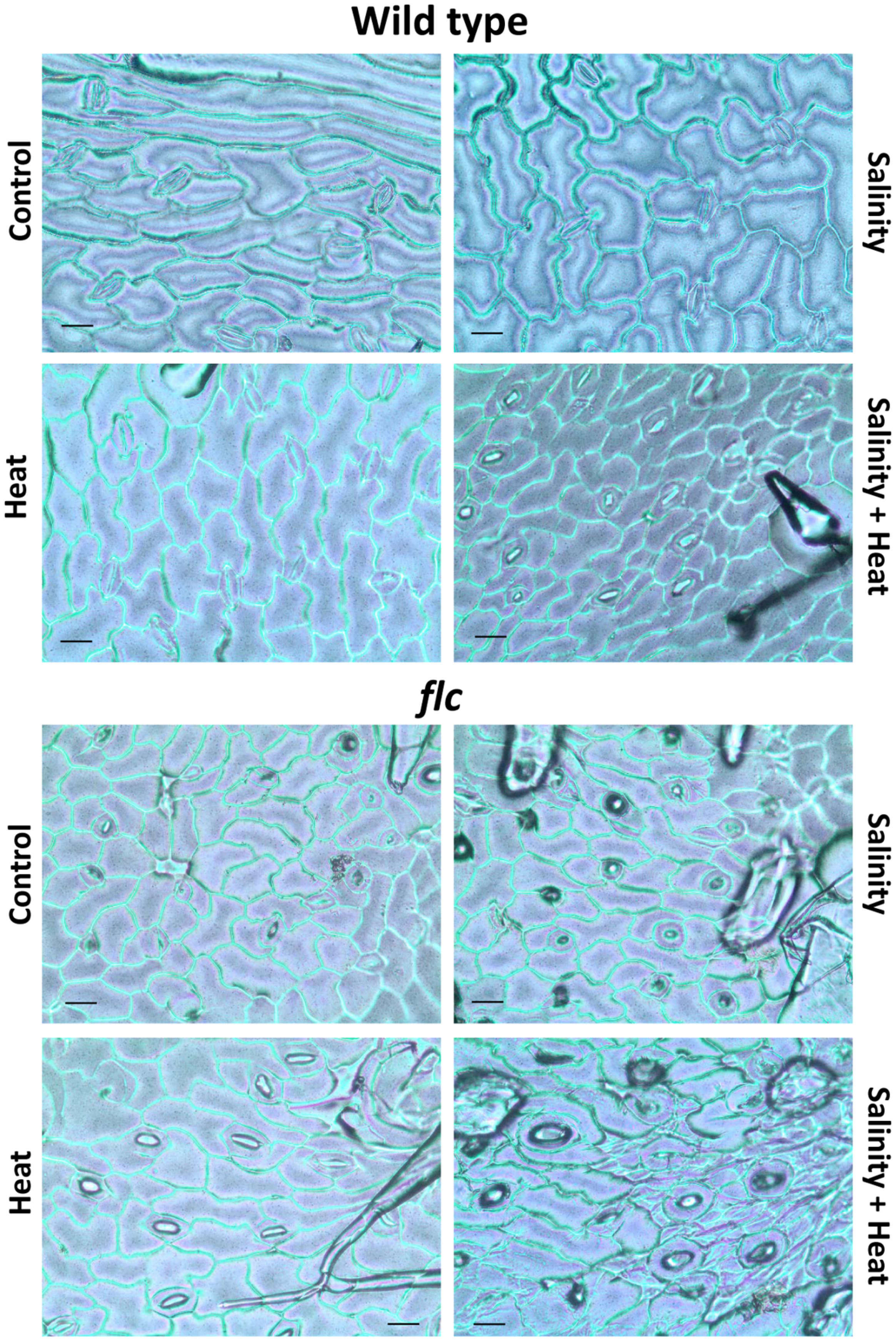


**Figure S1.** Light microscopy micrograph of adaxial leaf surface of tomato Wt and *flc* mutants grown under control, salinity, heat stress, or their combination. Scale bars: 100 µm
