## Supplementary material for "Specific ABA-independent tomato transcriptome reprogramming under abiotic stress combination": Figure S2

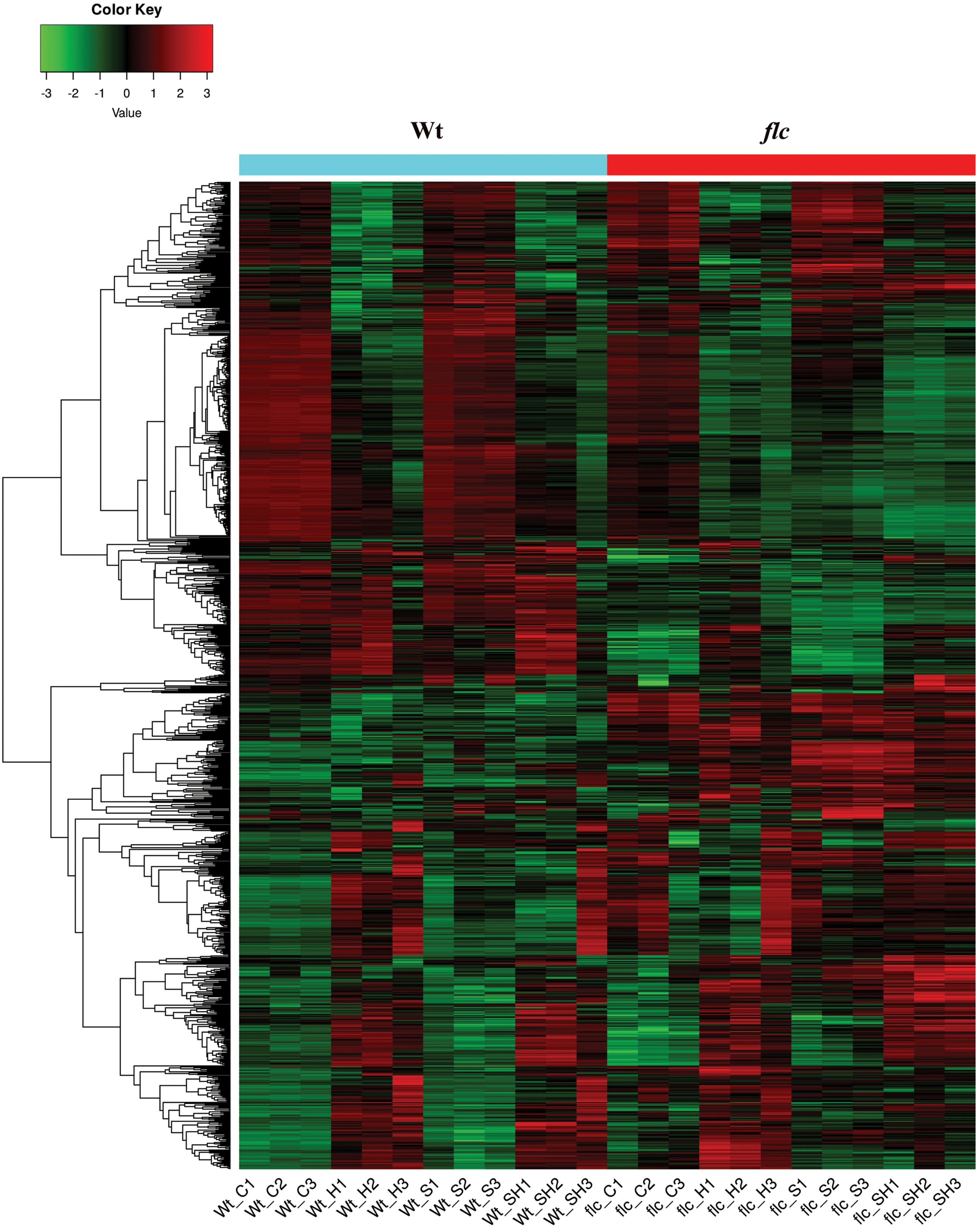


**Figure S2.** Heatmap for the gene expression obtained for each sample and genotype under every stress condition applied. C: Control; S: Salinity; H:Heat; SH: Salinity+Heat. 1, 2, and 3 represent the respective biological replication.
